# Dengue Virus Serotype 2 Infection Transcriptionally Reprograms Host Neutrophils to Establish an Immunosuppressive Microenvironment in Mice

**DOI:** 10.64898/2026.10.29.762704

**Authors:** Surender Rawat, Abhigya Chhetri, Sakshi Nimesh, Abhay Deep Pandey, Srikanth Sadhu, Gazala Siddqui, Sweety Samal, Amit Awasthi, Arup Banerjee

## Abstract

Dengue virus (DV) infection triggers profound alterations in host immune responses, yet the transcriptomic reprogramming of innate immune cells during disease progression remains incompletely understood. Here, we demonstrated that Dengue virus serotype 2 (DV-2) infection transcriptionally reprograms host neutrophils, establishing an immunosuppressive microenvironment that facilitates pathogenesis. Through scRNA transcriptomic profiling of bone marrow and spleen cells from the DV-2-infected AG129 mouse model, we delineated a distinct neutrophil signature that shows dynamic shifts in immature and activated populations over the course of dengue infection. We also characterized a specific Ly6G^low^ subpopulation, notably Fibrinogen-like protein 2 (FGL2)-expressing subsets, that appeared with disease progression. Serum FGL2 levels were significantly elevated in both DV-2-infected mice and human dengue-positive patients. Furthermore, treatment of splenocytes with FGL2 or FGL2-enriched neutrophil secretomes suppressed T cell functional activity and promoted an immunosuppressive phenotype. Together, these findings reveal how DV-2-driven neutrophil reprogramming shapes an immunosuppressive network, highlighting the importance of studying FGL2-expressing neutrophil subsets to understand dengue immunopathogenesis.

**Added value of this study:** This study provides the single-cell transcriptomic map of neutrophil heterogeneity during dengue virus serotype 2 infection in a mouse model, uncovering a critical temporal shift from early protective subsets to a severe-stage, immunosuppressive Ly6G^low^ and immature subpopulation. Progression triggers an abnormal cell subpopulation (Ly6G^low^) that secretes the soluble protein Fibrinogen-like protein 2 (FGL2) and suppresses T cell functional activity, promoting an immunosuppressive phenotype. By validating elevated plasma FGL2 levels in human dengue patients, this work identifies Ly6G^low^ neutrophil subsets that secrete FGL2 as a possible mediator of immunosuppression in severe dengue in the mouse model.

## Introduction

Neutrophils represent the most abundant and critical phagocytic cell population of circulating leukocytes, serving as the primary innate defence against systemic infection. In response to specific pathogens and environmental stimuli, these cells execute diverse effector functions, including phagocytosis, degranulation of proinflammatory enzymes, and the formation of neutrophil extracellular traps (NETs), and modulate adaptive immune responses (1–3). Neutrophils originate from hematopoietic stem cells (HSCs) in the bone marrow via granulopoiesis. The maturation process occurs sequentially. This sequence—progressing from myeloblasts to mature granulocytes—is defined by progressive reductions in cell size, nuclear segmentation, and the synthesis of cytoplasmic granules (1). While neutrophils have traditionally been viewed as terminally differentiated, recent advances in single-cell RNA sequencing (scRNA-seq) have revealed significant phenotypic and functional heterogeneity within neutrophil populations (4–8).

Under steady-state conditions, scRNA-seq has identified three distinct transcriptional subsets in circulation: immature neutrophils, functionally active mature neutrophils, and a specialised subset defined by interferon-stimulated genes (ISGs) (6, 9). However, during infection, granulocyte colony-stimulating factor (G-CSF) drives emergency granulopoiesis to meet increased demand (10, 11). This accelerated production often yields phenotypically and functionally altered neutrophils that can influence disease progression. Such heterogeneity is well-documented across various pathologies, including autoimmune disorders, oncology, and microbial infections (12–14). Consequently, neutrophils can adopt a broad spectrum of functional profiles, ranging from pro-inflammatory to immunosuppressive, depending on the microenvironmental context. In sepsis, the recruitment of IL1R2^+^ immature neutrophils is associated with worse patient outcomes (15). Emerging evidence suggests that dengue virus (DV) infection significantly affects neutrophil biogenesis and effector functions.

We previously demonstrated that dengue virus serotype 2 (DV-2) accelerates the differentiation of myeloid progenitor cells, leading to an increased systemic release of immature or newly formed neutrophils (16). During acute infection, neutrophils exhibit a highly activated phenotype (marked by increased surface expression of CD66b and CD16^bright^/CD62L^dim^) and generate neutrophil extracellular traps (NETs), which are implicated in vascular leakage and increased disease severity (16, 17). This excessive activation drives the inflammatory damage characteristic of severe dengue and other viral pathologies, positioning neutrophils as both critical pathogenic factors and viable therapeutic targets (18, 19).

Despite these important findings, research on neutrophils in dengue has been limited due to their short lifespan and the difficulty of handling them under experimental conditions. However, the emergence of single-cell RNA sequencing (scRNA-seq) has enabled the characterisation of neutrophil heterogeneity across diverse pathological states. A consensus is now forming around the use of scRNA-seq to identify specific disease-associated neutrophil phenotypes and their secretome for targetable therapy (19). Although functional heterogeneity between neutrophils and viral infection has been reported in the literature, there is a clear lack of information on whether dengue virus-induced granulopoiesis alters neutrophil phenotypes. Are those neutrophils functionally distinct, and do they influence disease outcome?

This study investigates neutrophil-mediated immune dysregulation during early and late-stage dengue progression using AG129 mice as a model for severe dengue. Using an unbiased scRNA-seq approach, we analysed neutrophil subsets in the bone marrow and spleen of DV-2-infected mice at 4 and 8 days post-infection (dpi), corresponding to mild and severe dengue, respectively. In addition, we explored mechanisms by which neutrophil subpopulations modulate T cell function. Our findings elucidate the transcriptomic landscape of dengue-associated neutrophils and highlight specific subpopulations associated with disease progression.

## Methods

### Ethics statement

The study adhered to ethical standards and obtained the necessary approvals for animal and human research. Animal procedures (RCB/IAEC/2021/112) involving C57BL/6 and AG129 mice were conducted at the NCR Biotech Science Cluster in Faridabad. Animals were maintained in accordance with the guidelines of the Committee for Control and Supervision of Experiments on Animals, Ministry of Environment and Forestry, Government of India. They were kept under a 12 h light/dark cycle with constant temperature and humidity. Food and water supply were *ad libitum*. Human blood experiments (RCB-IEC-H-25) were conducted at the Regional Centre for Biotechnology using previously collected stored samples, with institutional approval. Dengue patients’ (n= 12, fever cases >5 days, DENV IgM-positive, PCR-positive, Platelet count >50000) serum was collected and stored at −80 °C until use. Detailed information on screening and enrolment of dengue patients, as well as on blood sample collection, was provided earlier (20).

### Virus generation and titration

Dengue virus serotype 2 (DV-2, IND/P23085/1960) was propagated in C6/36 cells for *in vitro* use, while the mouse-adapted NR-15247 strain was used for animal experiments, with titers determined by foci-forming assay as described earlier (16, 21).

### AG129 mice and DV-2 infection

AG129 mice (6–8 weeks old, female, body weight 16-18g), maintained at the Experimental Animal Facility (EAF), NCR Biotech Science Cluster, were divided into two groups: mock and infected. Food and water were supplied ad libitum in the cages. Six animals (*n* = 6) in each group received 2 × 10^7^ Focus-Forming Units (FFU) of DV-2 intravenously. The mock group received an equal volume of PBS. From day 3 post-infection onwards, animals developed mild symptoms on days 6-8 and started showing severe symptoms. To capture immune cell status in bone marrow and spleen in mild and severe conditions, animals were killed at days 3-4 and 6-8 (depending on when symptoms appeared) using ketamine to minimise pain and distress during perfusion. For *ex vivo* studies, bone marrow cells were harvested from 7–8-week-old female mice. Subjects were monitored daily for weight changes and clinical symptoms, scored on a 1–5 scale ranging from piloerection to whole-body tremors. Following retro-orbital blood collection, mice were sacrificed at various time points to harvest the spleen, liver, and bone marrow.

### Isolation of cells from mock and DV-2-infected mice

Bone marrow (BM) cells were harvested from mouse femurs and tibias by flushing with PBS (2% FBS, 1 mM EDTA), then centrifuged and resuspended at 10^8^ cells/mL. Myeloid populations were identified via flow cytometry using CD11b, Ly6C, and Ly6G staining, while a subset was preserved in TRIzol for RNA isolation. Similarly, splenocytes were isolated using a cell strainer and stained for CD3, CD4, and CD8 to analyse and sort T cell populations. Detailed information about antibodies was provided in Table 1.

**Table 1:**
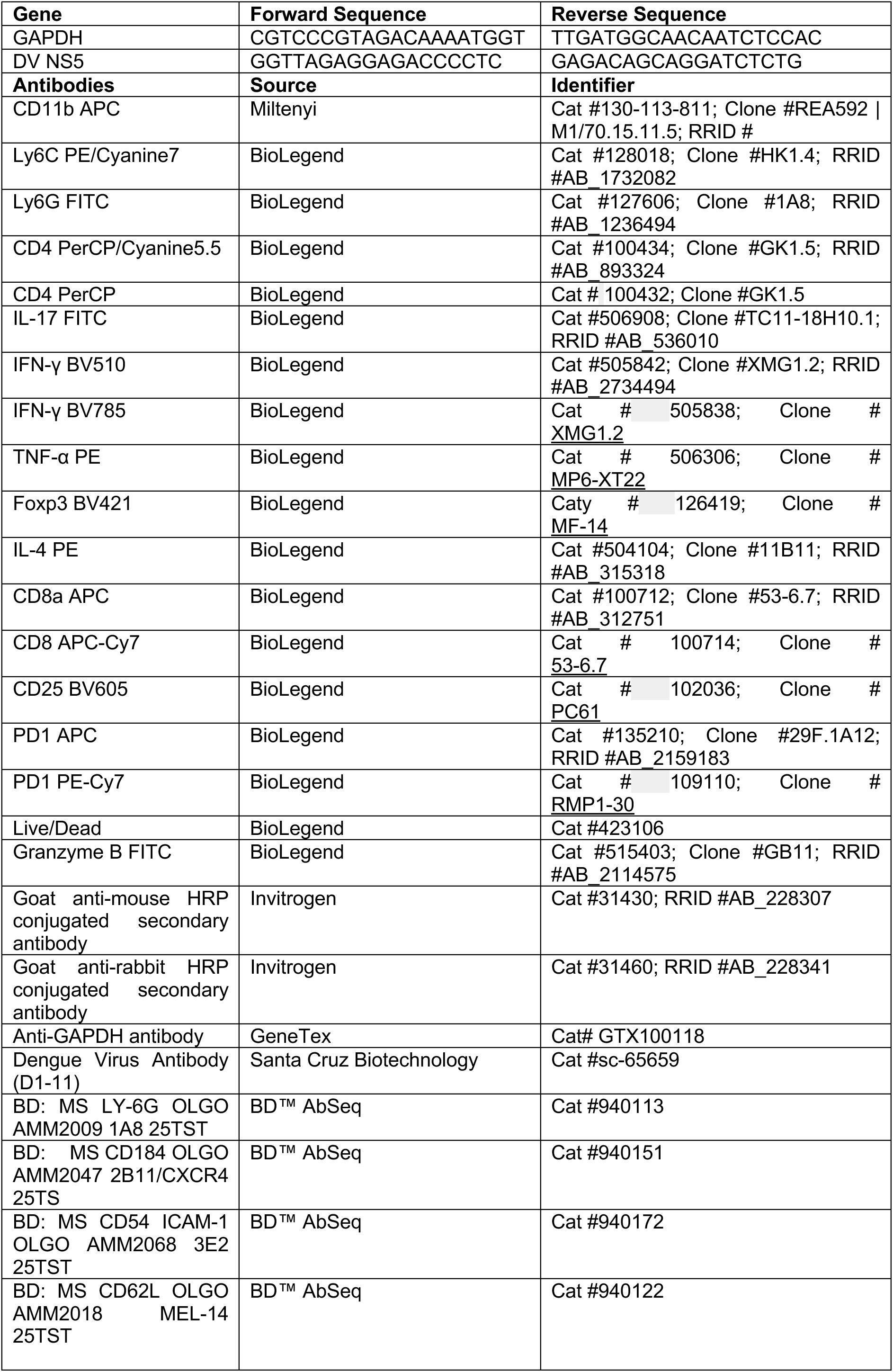
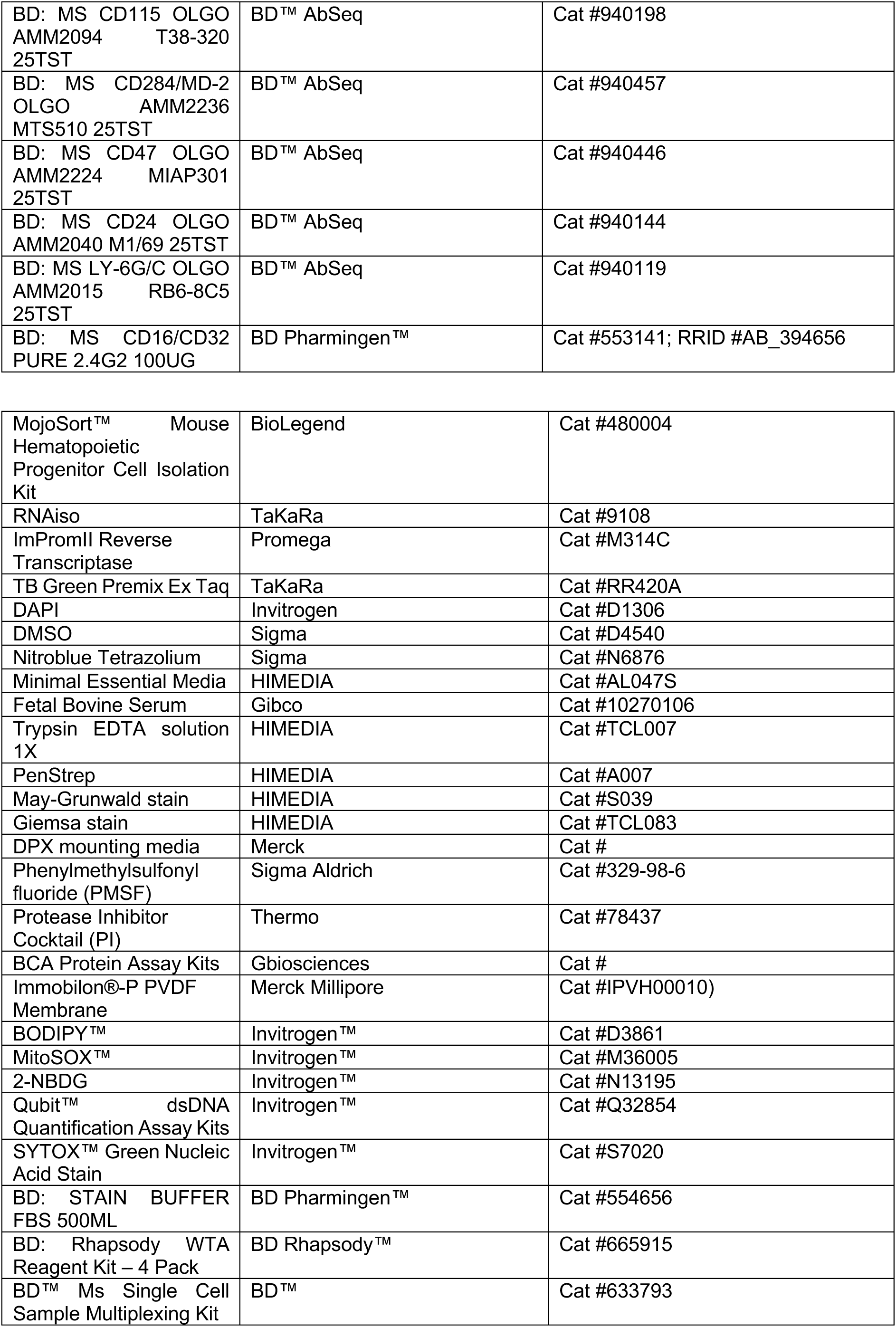
Primers, Antibodies and reagents list with RRID.

### BrdU proliferation assay

The BrdU proliferation assay was performed to assess the effect of dengue virus on bone marrow cell proliferation on days 3 and 6 post-infection with DV-2, according to the manufacturer’s guidelines (BD Pharmingen™ BrdU Flow Kits, Cat# 559619). For in vivo BrdU labelling, the mice were injected with 200 µL of BrdU solution (10mg/mL) via the intraperitoneal route for 4 hours. The BrdU-labelled mock and DV-2-infected mice were harvested at different time points, and bone marrow was collected. BrdU^+^ cells were quantified and analysed in flow cytometry (22) as described earlier.

### Colony formation assay

The colony formation assay was performed in semi-solid methylcellulose media under cytokine stimulation (R&D Systems, Cat# HSC007). Briefly, 2.5 × 10^4^ bone marrow cells (in 450 µL IMDM media) from mock and DV-2 infected mice after 3 and 6 days post-infection were mixed with 2.1 mL of methylcellulose media and allowed to settle for 20 minutes to remove air bubbles. Using a 16-gauge needle, 1.1 mL of the suspension was plated into each well or dish. Finally, the samples were incubated at 37 °C in a humidified CO_2_ incubator for 13–14 days without disturbance to allow for colony formation. Colonies were counted and characterised using the methods described earlier (23).

### Sorting of immune cells isolated from mice

To sort Ly6G^hi^ and Ly6G^low^ populations, bone marrow from mock and DV-2-infected mice was harvested six days post-infection. Cells were stained with CD11b, Ly6G, and Ly6C antibodies, then sorted using a BD FACS Aria Fusion. Following centrifugation, cells were used for co-culture and RNA isolation; pellets were stored in TRIzol at −80 °C. Similarly, CD4 and CD8 T cells were isolated from the spleens of AG129 mice mock-infected or DV-2-infected. Spleens were processed through a 70 µm strainer, stained, and sorted for use in co-culture and colony formation assays.

### T cells and neutrophil co-culture assay

The effects of the Ly6G^low^ and Ly6G^hi^ populations on the activation of CD8 T cells and distinct CD4 T cell subsets were analysed using a co-culture assay. Briefly, the sorted Ly6G^low^ and Ly6G^hi^ populations from bone marrow cells were co-cultured with purified CD4 and CD8 T cells isolated from splenocytes. The CD4 and CD8 T cells were cultured in a 96-well plate pre-coated with anti-CD3 antibody and soluble anti-CD28 antibody for expansion, and the Ly6G^low^ and Ly6G^hi^ populations were added at a 1:10 ratio and incubated for 3 days at 37 °C in a CO_2_ incubator. The cells were then activated with PMA (50 ng/mL; Sigma-Aldrich) and Ionomycin (1.0 µg/mL; Sigma-Aldrich) in the presence of Monensin (BD Biosciences), followed by staining for surface markers and intracellular cytokines.

### Intracellular and surface staining of CD4 and CD8 T cells

Intracellular staining was performed by incubating the cells in perm/wash buffer, followed by staining with antibodies for 30 min at 4 °C. The antibodies included anti-mouse CD4 (PercCP/Cyanine5.5; Clone GK1.5), anti-mouse IL-17 (FITC; Clone TC11-18H10.1), anti-mouse IL-10 (APC; Clone JES5-16E3), anti-mouse IL-4 (PE; Clone-11B11), and anti-mouse IFN-γ (BV510; Clone XMG1.2) for staining different subsets of CD4 T cells, while CD8 cells were stained with anti-mouse CD8a (APC; Clone 53-6.7), anti-mouse PD1 (APC; Clone 29F.1A12), and anti-mouse Granzyme B (FITC; Clone GB11) antibodies. After incubation, the cells were washed with PBS, and the samples were acquired on a FACS Symphony^TM^ instrument (BD Biosciences), using BD FACS Suite software version 1.0.6 (BD Biosciences, San Jose, CA, USA), and analysed with FlowJo software version VX (Tree Star, San Carlos, CA, USA). Detailed information about antibodies was provided in Table 1.

### Quantification of dsDNA using the PicoGreen assay

The Quant-iT PicoGreen dsDNA Reagent, an ultrasensitive fluorescent nucleic acid stain, was used to quantify double-stranded DNA (dsDNA) in plasma samples from mock- and infected-mice according to the manufacturer’s instructions. Briefly, a standard curve was made by serially diluting (tenfold) the 2 mg/mL stock solution of λ DNA in Tris-EDTA (TE) buffer. The working solution of the quant-iT reagent was made by diluting 50 µL of the reagent in 9950 µl of TE buffer (1:199). 50 µL of plasma samples and the standards were added to 150 µL of the working solution of the quant-iT reagent, and the mixture was incubated for 5 minutes. The samples were then analysed by measuring absorbance using a SpectraMax i3x plate reader (Molecular Devices, USA) at an Excitation/emission wavelength of 488/520 nm.

### Quantitative real-time (qPCR) assay

Total RNA was isolated using RNAiso (TaKaRa, Dalian, China), and RNA concentration was measured using a NanoDrop 1000 spectrophotometer. Reverse transcription (RT) was then performed with 1 µg total RNA using ImPromII Reverse Transcriptase (Promega). cDNA was subjected to real-time PCR using TB Green Premix Ex Taq (Tli RNase H Plus) (TaKaRa) and Real-time PCR System Quant Studio 6 Flex (Applied Biosystems). The qRT-PCR primer sequences used are listed in Table 1.

### Western blotting

Tissue samples were homogenised and lysed in the lysis buffer containing 20 mM Hepes pH 7.5, 10 mM EGTA, 1% NP40, 2.5 mM MgCl_2_, 40 mM β-Glycerophosphate, 2 mM NaV, 1 mM DTT, 1 mM PMSF, and 1x Protease Inhibitor cocktail (Thermo Fisher Scientific, Waltham, MA, USA) for 1 h on ice. The lysates were cleared by centrifugation, then boiled with 6X Laemmli buffer at 95 °C for 10 min, and equal amounts of protein were loaded, separated by SDS-PAGE, and transferred to a PVDF membrane. The membranes were blocked with 5% BSA for 1 h at room temperature and incubated with primary antibodies against Cas9 and GAPDH (GeneTex) in 1% BSA overnight at 4 °C. The following day, the membranes were incubated with the HRP-conjugated secondary antibody for 1 h at room temperature, then visualised using ImageQuant (LAS4000).

### 2-NBDG uptake assay

Cells were washed with 1XPBS twice and allowed to starve at 37 °C for 1 h in glucose-free RPMI media. After starvation, 150 µM 2-NBDG (2-(N-(7-Nitrobenz-2-oxa-1,3-diazol-4-yl) Amino)-2-Deoxyglucose was added to cells, and incubation was done for another 25 minutes at 37 °C. After staining, washes were given, and data were collected using a FACSVerse Flow Cytometer (BD Biosciences).

### MitoSox staining

Cells were washed twice with 1× PBS, and MitoSox (1 μM) was added for 15 minutes at 37 °C. After staining, washes were performed, and data were collected using a FACSVerse Flow Cytometer (BD Biosciences).

### Bodypi staining

Cells were washed twice with 1× PBS, and Bodypi (2 μM) was added for 20 minutes at 37 °C. After staining, washes were performed, and data were collected using a FACSVerse Flow Cytometer (BD Biosciences).

### Giemsa staining

Sorted cells (0.02 × 10^6^) from bone marrow were cytospun at 500g for 5 min onto slides. May-Grunwald staining was performed for 20 min, followed by washes and counterstaining with Giemsa Stain (1:10 dilution) for 30 min. The slides were washed, dried, and mounted with DPX mounting media. Imaging was done using a Nikon Eclipse TS2 microscope with a 60× objective.

### Immunofluorescent confocal microscopy

Sorted cells were seeded at a density of 0.2 × 10^6^ cells per well in serum-free RPMI media onto round glass coverslips coated with poly-L-Lysine in a 24-well plate. The cells were allowed to adhere for an hour. After 3 h of incubation, the cells were fixed with 4% paraformaldehyde and permeabilised with 0.1% Triton X-100. For nuclear visualisation, cells were stained with DAPI (5 μg/mL). Cells were mounted onto slides with mounting media (Prolong Gold, Invitrogen) and observed using a Confocal DMI 6000 TCS-SP8 microscope (Leica Microsystems, Wetzlar, Germany).

### Extracellular DNA release using Sytox fluorescent

For quantification of extracellular DNA release, cells were seeded at a density of 0.2 × 10 ^6^ cells per well in a 96-well plate. The cells were treated with PMA (100ng/ml) and incubated for 30 minutes. Cell culture supernatant was removed, and Sytox green was added at a final concentration of 0.25 µM for 15 minutes. The absorbance was measured using a SpectraMax i3x plate reader (Molecular Devices, USA).

### Single-cell RNA sequencing

Bone marrow and spleen cells from mock-infected (n=3) and infected (n=3) mice were isolated, filtered, and assessed for >90% viability using trypan blue. All steps were performed at 4°C.

Samples (0.2–0.3 million cells each) were labelled with BD sample tags, incubated on ice, and pooled. Following AbSeq staining, approximately 1 × 10^6^ pooled cells were loaded onto a BD Rhapsody Cartridge for single-cell capture. After adding magnetic capture beads and washing, cells were lysed, and beads were recovered for reverse transcription with target-specific primers. Finally, WTA, AbSeq, and Sample Tag sequencing libraries were prepared, with quality and concentration verified using Qubit and Bioanalyzer TapeStation before sequencing.

### Preprocessing and bioinformatics analysis of single-cell RNA-seq data

Raw single-cell RNA-seq data generated using the BD Rhapsody platform were processed on the Seven Bridges platform (Available online at: https://www.sevenbridges.com/platform/) to generate gene-expression, AbSeq, and sample-tag matrices. Reads were aligned to the Mus musculus mm10 reference genome and quantified using unique molecular identifiers (UMIs). Downstream analysis was performed in R version 4.6.0 using Seurat version 5.5.1. Bone marrow and spleen samples from mock- and DV-2-infected mice at 4 and 8 days post-infection (dpi) were analysed.

Cells were retained if they contained 200–6000 detected genes, 500–50,000 RNA counts, and ≤20% mitochondrial reads. Multiplets identified using the BD sample tags were excluded. Data were normalised using SCTransform with glmGamPoi version 1.24.0 and vst.flavor = “v2”. Principal component analysis (PCA) was performed on the normalised data. A shared-nearest-neighbour graph was constructed using the first 25 principal components, followed by graph-based clustering in Seurat at a resolution of 0.5. t-SNE was generated using the first 25 principal components. UMAP was generated from the PCA space using the first 25 principal components, cosine distance, and a fixed random seed of 1234. Bone marrow and spleen cells were analysed jointly, and no explicit batch or tissue integration method was applied. Major cell populations were annotated using established lineage markers. The original T-cell population was further separated into CD4 and CD8 T cells using PCA and graph-based clustering with the first 15 principal components and a resolution of 0.05. Cluster marker genes were identified using Seurat FindAllMarkers with positive markers only, a minimum expression fraction of 0.10, and a log-fold-change threshold of 0.25.

Granulocytic populations were assigned to developmental and functional states G0– G5c using marker expression and published neutrophil signatures, including those described by Xie et al. (6). CMP and GMP populations were grouped as G0. G1–G3 represented progenitor and immature neutrophil states, G4 represented mature/resting neutrophils, G5a represented an immunosuppressive state, G5b represented an interferon-responsive ISG^+^ state, and G5c represented an activated/inflammatory state. Existing cell populations were retained, and no additional neutrophil re-clustering was performed. State assignments were checked using marker-gene expression, dot plots, heatmaps, tissue distribution, and temporal distribution.

Functional properties of the granulocytic states were assessed using gene signatures for azurophil, specific, gelatinase, and secretory granules, as well as NADPH oxidase activity, chemotaxis, and phagocytosis (6). For each cell, the functional score was calculated as the mean log-normalised RNA expression of the genes detected in each signature. Chemotaxis and phagocytosis genes were obtained from mouse Gene Ontology annotations GO:0030593 and GO:0006911, respectively. The NADPH oxidase signature included Cyba, Cybb, Ncf1, Ncf2, Ncf4, Rac1, and Rac2. Functional scores were compared across granulocytic states, tissues, and infection time points. Granulocyte maturation was further analysed using Slingshot (24). RNA expression was log-normalised, variable genes were selected, and PCA was recalculated to infer trajectories. All granulocytic states from G0 to G5c were included. G0 was used as the starting state when present. Pseudotime was calculated along the inferred lineage, while the same UMAP coordinates were retained for visualisation. Normalised Ly6g expression was plotted along pseudotime and visualised using a LOESS-smoothed curve.

Recovered cell numbers and relative proportions were calculated for each tissue and time point. These values represent cells recovered after sequencing and quality control and were not treated as absolute tissue cell counts. Since each tissue and time-point condition was represented by one scRNA-seq sample, cell-level differences in abundance and functional scores were treated as descriptive and not used in biological-replicate statistical tests.

### Human neutrophil isolation

Blood was collected from healthy, consenting donors in 6-mL EDTA-coated vials. The neutrophil was then collected using the EasySep™ Direct Human kit. Neutrophil Isolation Kit (Cat# 19666, StemCell Technologies, Canada) using the negative selection principle. The antibody cocktail and the rapid sphere were added to the blood and incubated for 5 min each. After incubation, 1X DPBS was used to dilute the blood, and the tubes were kept in the EasySep™ Magnet for 5min. The steps were repeated at least 3 times to obtain purified neutrophils. The suspension was centrifuged at 1000 rpm for 10 min. The obtained pellet was washed with 1x DPBS. The cells were resuspended in incomplete RPMI-1640, counted using a hemocytometer, and then used for experiments.

### FGL2 ELISA

Purified neutrophils were seeded in 12-well plates at a density of 1 × 10⁶ cells per well, infected with DV-2 (MOI 1), and incubated for 12 and 24 hours; culture supernatants were collected. For serum analysis, mouse and human samples were diluted 1:50 in the kit-provided assay buffer before ELISA. Sorted Ly6G^low^ cells were seeded in 24-well plates at a density of 0.5 × 10⁶ cells per well and infected with DV-2 (MOI 1) for 24 hours, followed by the collection of supernatants. Levels of FGL2 in culture supernatants and serum samples were quantified using Mouse FGL2 (Fibrinogen-like Protein 2) ELISA Kit (Cat #EM1024) and Human FGL2 ELISA Kit (Cat #EH3064) (Finetest, Boulder, CO, US), according to the manufacturer’s instructions.

### FGL2 T-cells interaction study

Single cell suspensions were made from the spleen and lymph nodes of 6–8 week-old mice. To investigate the immunomodulatory effect of FGL2 on T-cell responses, splenocytes were cultured in a plate pre-coated with anti-CD3 antibody (2.0 µg ml^−1^, 145-2C11, BioXcell). Cells were maintained either in the absence of additional stimulation, in the presence of soluble anti-CD28 (2.0 µg /mL, PV1, BioXcell) to provide co-stimulatory signalling, or in the presence of recombinant FGL2 protein (1 µg/mL), either alone or in combination with anti-CD3/CD28 stimulation. Cultures were maintained at 37 °C and 5% CO₂ for 48 hours to allow for T-cell activation and functional modulation (25).

To evaluate the effects of neutrophil-derived factors under viral conditions, Ly6G^low^ and Ly6G^hi^ cell populations from mock- and dengue-infected cells were cultured. The resulting conditioned supernatants were collected for stimulation assays. Draining lymph node cells were cultured in conditioned media comprising supernatants from both uninfected and dengue-infected Ly6G^low^ and Ly6G^hi^ populations, with corresponding mock controls. Cells were incubated for 48 hours under standard culture conditions to assess the impact of FGL2 on T-cell responses. For intracellular cytokine analysis, cells were harvested, washed, and stained with a viability dye Zombi NIR to exclude dead cells. Surface staining was performed using fluorochrome-conjugated antibodies against CD4 (Percp; 1:1000), CD8 (APC-Cy7; 1:1000), CD25 (BV605; 1:700) and PD-1 (PE-Cy7; 1:1000). Cells were subsequently fixed and permeabilized using a BD Cytofix/permeabilization kit (#554714, BD Biosciences), followed by intracellular staining for IFN-γ (BV785; 1:400), TNF-α (PE; 1:400), perforin (APC; 1: 500), IL-10 (BV650; 1: 400), and Foxp3 (BV421; 1:200). Data acquisition was performed using a Cytek Northern Lights™ flow cytometer, and analysis was carried out using FlowJo software (Tree star)(26, 27).

### Statistical analysis

All statistical analyses for scRNA-seq data were performed with the R software package. The significance of DEGs and transcripts was analyzed and a *P*-value of <0.05 and Log2 fold change (LFC) value ≥ 0. 5 was considered to indicate a significant difference. Data were analysed in GraphPad Prism 8.0 and presented as Mean ± SD of at least three independent experiments with two replicates. Data were analyzed for Gaussian distribution. Statistical significance was determined either with a two-tailed Student’s *t*-test or one-way ANOVA followed by Tukey’s multiple comparison tests when three or more groups were analyzed. No test for outliers was performed. A p-value of <0.05 was considered statistically significant.

### Data deposition and materials sharing

#### Code availability

All custom code used for scRNA-seq analysis is publicly available on GitHub at https://github.com/abpseq/Dengue-scRNAseq-analysis

#### Data Availability

Raw and processed sequencing data have been deposited in the Gene Expression Omnibus (GEO) (GSE329847) database. The data will be accessible upon publication.

## Results

### DV-2 infection of AG129 mice develops manifestations associated with severe dengue and infects the bone marrow progenitor cells

To elucidate neutrophil dynamics in dengue and translate single-cell insights into clinical applications, we utilised a previously established mouse model for DV-2 infection (21). This model effectively recapitulates severe human dengue, including increased vascular permeability and lethal disease (21, 28–30). Six-to eight-week-old AG129 mice were intravenously challenged with the mouse-adapted DV-2 strain NR-1524 and monitored daily for morbidity (Fig. 1A) (21). Infected mice displayed progressive clinical deterioration, including ruffled fur, hunched posture, and hind limb paralysis by days 6–7, reaching a moribund state by days 8–10 post-infection (pi) (Fig. 1 B). Significant weight loss was observed from day 4 pi (Fig. 1C), with 100% mortality occurring by days 10–12 pi (Fig. 1D-E). We sacrificed the mice on days 3-4 and 6-8 post-injection and harvested blood, liver, spleen, and BM for different assays. The infected mice showed significant splenomegaly at 8 dpi (Fig. 1F). The total WBC count increased significantly at 4 dpi compared to mock controls. This trend continues upward through 8 dpi. There was a slight, non-significant (ns) increase in lymphocytes at 4 dpi, followed by a more substantial increase by 8 dpi. A statistically significant (*p< 0.05) increase in neutrophil count occurs early (4 dpi) and persists through 8 dpi, suggesting an active innate immune response and potential inflammation (Fig. 1G). Additionally, infected mice exhibited symptoms of severe dengue, including thrombocytopenia (Fig. 1H) and significant vascular leakage in multiple organs, including the spleen, liver, kidney, and intestine (Fig. 1I).

**Fig. 1.**
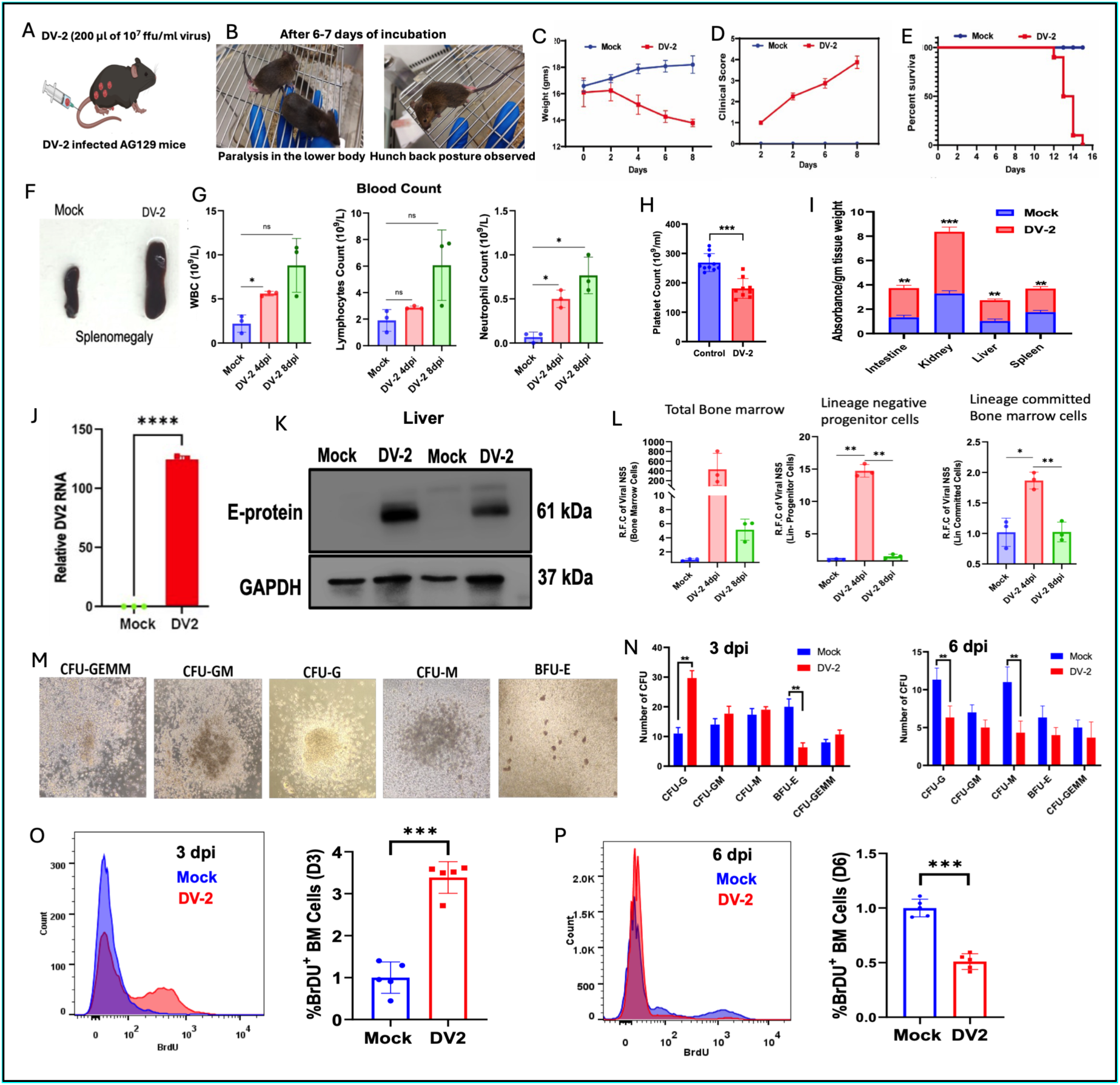
Characterization of Dengue virus 2 infection dynamics, pathology, and hematopoietic alterations in AG129 mice. **(A)** Experimental scheme of Dengue virus 2 (DV-2) infection (200 ul of 10^7^ ffu/ml) in AG129 mice. **(B)** Representative photographs showing clinical manifestations, including lower body paralysis and a hunchback posture, observed in DV-2–infected mice 6–7 days post-incubation. **(C-E)** Longitudinal monitoring of body weight in grams (**C**), clinical scores (**D**), and Kaplan-Meier survival curves (**E**) comparing Mock- and DV-2–infected AG129 mice over time. **(F)** Representative image displaying splenomegaly in DV-2–infected mice compared to Mock controls. **(G)** Complete blood counts detailing total white blood cell (WBC), lymphocyte, and neutrophil concentrations (10^9^ mL) in peripheral blood across Mock, 4 days post-infection (4 dpi), and 8 dpi groups. **(H)** Platelet counts (10^9^ ml) in control and DV-2–infected mice. **(I)** Vascular leakage evaluation quantified by Evans blue dye absorbance per gram of tissue weight across intestine, kidney, liver, and spleen in Mock- and DV-2–infected groups. **(J)** Relative DV-2 RNA expression in Mock versus DV-2 conditions measured by RT-qPCR. **(K)** Immunoblot detection of viral Envelope (E)-protein (61kDa) in liver homogenates from Mock- and DV-2–infected mice. GAPDH 37kDa) served as a loading control. **(I)** Relative fold change (R.F.C) of viral NS5 transcript levels in total bone marrow (left), lineage-negative (Lin-) progenitor cells (middle), and lineage-committed bone marrow cells (right) measured at Mock, 4 dpi, and 8 dpi. **(M)** Representative brightfield colony morphology images from colony-forming unit (CFU) assays, including CFU-GEMM, CFU-GM, CFU-G, CFU-M, and burst-forming unit-erythroid (BFU-E). **(N)** Quantification of total colony numbers per lineage at 3 dpi (left) and 6 dpi (right) following Mock or DV-2 challenge. **(O, P)** Representative flow cytometry histograms and bar graph quantifications of BrDU integration (BrDU^+^) in bone marrow cells at 3 dpi (**O**) and 6 dpi (**P**) in Mock-versus DV-2–infected mice. Data in line plots (**C**, **D**) and bar graphs (**G**–**J**, **L**, **N**–**P**) are presented as mean ± s.d. Statistical significance was calculated using two-sided Student’s t-test or one-way ANOVA with post-hoc test; ns, not significant; *P < 0.05,P < 0.01, ***P < 0.001, ****P < 0.0001

We isolated RNA from the spleens of infected mice. The qRT-PCR and immunoblot showed abundance of viral RNA in the infected spleen and liver, respectively (Fig. 1J, K). Dengue virus is known to infect bone marrow and hematopoietic cells (31, 32). In our experimental model, we observed elevated viral RNA levels in total bone marrow, lineage-negative progenitor cells, and lineage-committed cells, suggesting that the virus can replicate in the bone marrow cells specifically during early stages of infection (Fig. 1L). Given the detection of viral RNA in progenitor cells, we investigated the impact of DV-2 infection on hematopoietic stem and progenitor cell (HSPC) expansion in the bone marrow (BM). Using colony-forming assays (CFA) in methylcellulose medium, we assessed the proliferation and differentiation potential of single cells over 14 days (Fig. 1M). Assays performed at day 3 post-infection (3 dpi) revealed a significant increase in colony-forming unit-granulocyte (CFU-G) frequency, alongside a sharp decline in burst-forming unit-erythroid (BFU-E) colonies compared to mock-infected controls (Fig. 1M, N).

By day 6 pi, however, a marked reduction in both CFU-G and colony-forming unit-macrophage (CFU-M) numbers was observed in DV-2-infected mice (Fig. 1N). This decline in CFU-M suggests a depletion of megakaryocyte progenitors, which may contribute to the observed thrombocytopenia. Collectively, these data indicate that DV-2 infection induces an initial phase of HSPC hyperproliferation, followed by a hypoproliferative state in bone marrow.

To further assess the impact of DV-2 on bone marrow proliferation, we used the BrDU assay (Fig. 1O, P). This assay revealed that at earlier stages of infection, there’s a near threefold increase in %BrDU^+^ cells with a subsequent decrease in %BrDU^+^ cells at later stages of infection in DV-2 infected mice with respect to their mock-infected control (Fig. 1O, P), suggesting the direct infection of bone marrow progenitor cells at the initial stage of infection might cause the observed proliferative surge.

### Single-Cell Transcriptomic Analysis Reveals Altered Neutrophil Maturation Dynamics and Heterogeneity During Dengue Virus Infection

Building on our previous finding that dengue infection expands myeloid cells in murine bone marrow (16), we investigated whether the virus induces increased myelopoiesis, which may generate functionally altered myeloid cells and contribute to severe disease. To dissect cellular heterogeneity and dynamic responses during Dengue virus (DV-2) infection, bone marrow and spleen cells were harvested from mock and infected mice at 4 and 8 days post-infection (dpi) for single-cell RNA sequencing (Fig. 2A). Uniform Manifold Approximation and Projection (UMAP) analysis identified major immune cell lineages and tissue-specific distributions in the bone marrow and spleen (Fig. 2B, C) using established canonical markers. Tissue distribution of identified clusters was depicted in Fig. 2C. Additionally, to dissect neutrophil heterogeneity, we also linked the neutrophil population defined by our scRNA-seq to the data reported by (6), which categorised neutrophils based on the core signature, and did not change even under infection. Our targeted sub-clustering and functional program analysis of this granulocytic compartment identified distinct maturation and activation states, annotated chronologically from G0 to G5c. (Fig. 2B). Tissue distribution of cell clusters was depicted in Fig. 2C. Granulocytes were further sub-classified into discrete developmental and functional states, designated as G0 through G5c. These states encompass immature progenitors (G0–G3), mature resting neutrophils (G4), and specialized subsets including immunosuppressive (G5a), interferon-responsive (G5b, ISG^+^), and activated/inflammatory (G5c) populations (Fig. 2B, F). Analysis of maturity programs using correlation matrices revealed coordinated transitions among granulocytic states (Fig. 2D), with bubble plots confirming stage-specific marker gene expression profiles (Fig. 2E).

**Fig. 2.**
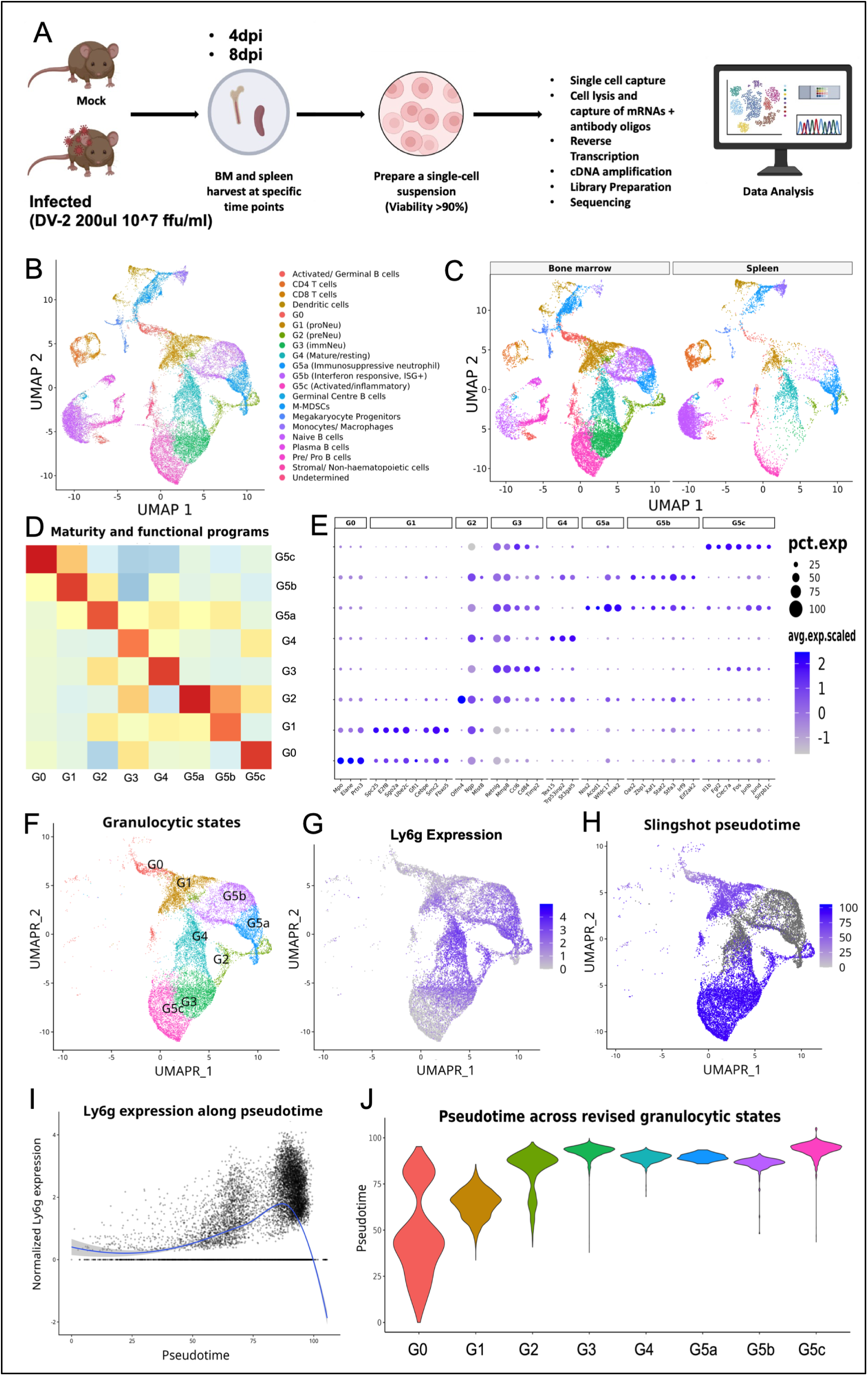
Single-cell RNA sequencing landscape and lineage trajectory of granulocytic states during dengue virus infection. **(A)** Schematic diagram of the experimental workflow. Mice were either mock-infected or infected with Dengue virus 2 (DV-2; 200 µl at 10^7 ffu/ml). Bone marrow (BM) and spleen tissues were harvested at 4- and 8-days post-infection (dpi), single-cell suspensions were prepared (viability > 90%), and single-cell RNA sequencing libraries were prepared for high-throughput sequencing and bioinformatic analysis. **(B)** UMAP plot showing major immune cell types identified across the combined single-cell dataset, including B cell, T cell, dendritic cell, myeloid, megakaryocyte/platelet progenitor, and stromal populations. **(C)** UMAP embedding of immune cell clusters split by tissue of origin: bone marrow (left) and spleen (right). **(D)** Correlation heatmap illustrating maturity and functional program similarities among defined granulocytic states (G0 to G5c). **(E)** Dot plot depicting marker gene expression profiles across individual granulocytic states (G0–G5c). Dot size represents the percentage of cells expressing a given gene (pct.exp), and color intensity indicates scaled average expression (avg.exp.scaled). **(F)** Sub-clustering UMAP visualization highlighting defined granulocytic developmental and functional states (G0, G1: proNeu, G2: preNeu, G3: ImmNeu, G4: Mature/resting, G5a: Immunosuppressive, G5b: ISG+, and G5c: Activated/inflammatory). **(G)** UMAP feature plot showing normalized *Ly6G* gene expression levels across granulocytic clusters. **(H)** Slingshot trajectory inference overlaid on the UMAP projection, coloured by pseudotime values that reflect developmental progression. **(I)** Line plot showing normalized *Ly6G* expression dynamics along inferred Slingshot pseudotime. The solid blue line represents a smoothed local regression (LOESS) fit. **(J)** Violin plots displaying the distribution of pseudotime scores across revised granulocytic states (G0 through G5c).

Furthermore, Slingshot pseudotime trajectory mapping tracked granulocyte differentiation from early progenitors (G0) toward mature and specialized terminal states (Fig. 2G-J). Interestingly, G5c neutrophils exhibited lower Ly6G expression than other mature populations (Fig. 2G). Pseudotime analysis computationally orders cells along a continuum based on gradual shifts in their gene expression profiles. The pseudotime score is calculated by measuring transcriptional similarity among cells and mapping them onto a continuous developmental path. A low G0 score indicates that the cell is in an early phase of the trajectory (Fig. 2H). Based on data obtained from ordered pseudo-time scoring across granulocytic populations, a clear hierarchy was established, originating at G0 (GMP) with a lower pseudo-time score and progressing through intermediate G1–G3 precursor stages toward mature resting (G4) and specialised activated/inflammatory subsets (G5a–G5c). This developmental progression correlated with an upregulation of Ly6G expression along pseudotime, validating the transition dynamics across granulocytic compartments during infection (Fig. 2I, J).

### Dynamics of Neutrophil States and Functional Scores Across Tissues During Dengue Virus Infection

After confirming the presence of different neutrophil subsets in the bone marrow and spleen, we examined their temporal appearance across time points. To evaluate tissue-specific shifts and temporal changes in myeloid and neutrophil populations during DV-2 infection, single-cell analyses were performed across bone marrow (stages ST1 to ST3 for Mock, 4 dpi, and 8 dpi, respectively) and spleen (stages ST4 to ST6 for Mock, 4 dpi, and 8 dpi, respectively).

Proportional compositions and cell counts across selected states (ST1–ST6) revealed major redistributions of neutrophil states in response to infection (Fig. 3A, B).

**Fig. 3.**
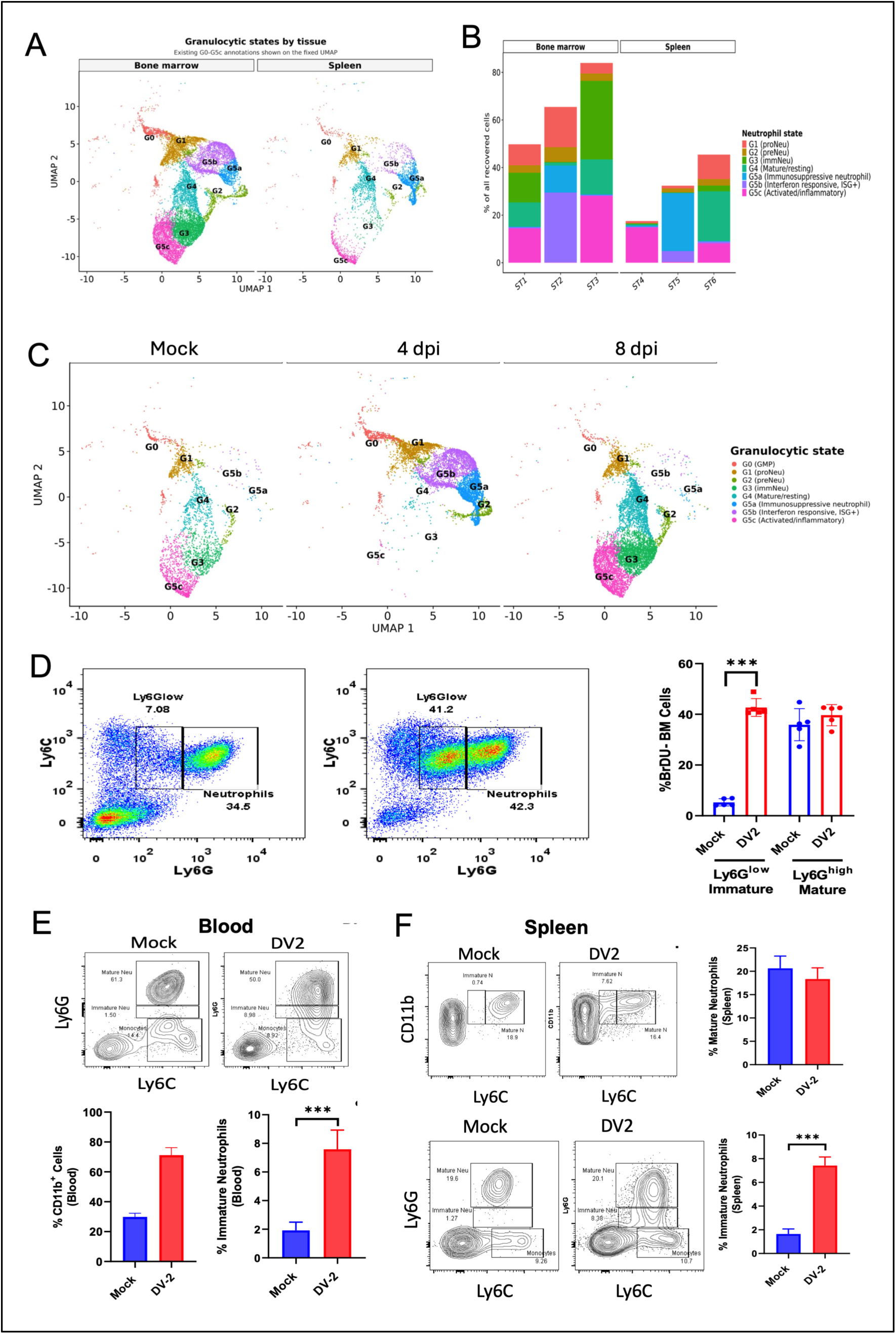
Single-cell landscape and developmental dynamics of neutrophil subsets during DV-2 infection. **(A)** UMAP visualization showing granulocytic states across bone marrow (left) and spleen (right) tissue microenvironments. Subsets are annotated based on developmental and functional stages (G0 to G5c). **(B)** Stacked bar plot illustrating the percentage composition of distinct neutrophil states (G1: proNeu, G2: preNeu, G3: ImmNeu, G4: Mature/resting, G5a: Immunosuppressive neutrophil, G5b: Interferon-responsive ISG+1, and G5c: Activated/inflammatory) across bone marrow and spleen tissue samples (ST1–ST6). **(C)** UMAP embedding depicting the distribution and temporal dynamics of granulocytic states under Mock, 4 days post-infection (4 dpi), and 8 dpi conditions following DV-2 challenge. **(D)** Representative flow cytometry plots (left) gating Ly6C versus Ly6G expression to identify Ly6G^low^ immature neutrophils and mature neutrophils in bone marrow, along with bar graph quantification (right) showing the percentage of BrDU^−^ cells in Ly6G^low^ immature and Ly6G^hi^ mature bone marrow (BM) populations following Mock or DV-2 infection. **(E)** Representative flow cytometry contour plots (top) and summary quantification (bottom) showing the proportions of total CD11b+ cells and immature neutrophils in peripheral blood of Mock- and DV-2-infected mice. **(F)** Representative flow cytometry contour plots (top and bottom-left) and bar graph quantification (right) showing the frequencies of mature and immature neutrophil populations in the spleen of Mock- and DV-2-infected mice. Data in bar graphs (**D**–**F**) are presented as mean ± s.d. Statistical significance was determined using two-sided Student’s t-test or one-way ANOVA; ***P < 0.001.

Quantitative breakdown of neutrophil states, expressed as a percentage of all recovered cells, reveals striking tissue-specific differences between the bone marrow and spleen. Bone marrow showed dynamic shifts in immature and activated populations over time, while splenic compartments exhibited distinct increases in Immunosuppressive (G5a) and interferon-responsive (G5b) subsets following infection (Fig. 3B).

UMAP visualizations categorized by tissue and infection time point (Mock, 4 dpi, and 8 dpi) demonstrated clear clustering of neutrophil states, including G0 through G3 progenitors, mature G4 resting cells, and specialized G5 subsets (G5a, G5b, and G5c). Infection induced marked spatial reconfiguration of these clusters in both bone marrow and spleen environments (Fig. 3A,C).

Under uninfected conditions, UMAP clustering showed a balanced distribution of baseline populations, with resting mature states (G4) and standard developmental progenitors (G0 through G3) dominating. Specialized subsets like G5a, G5b, and G5c maintained low baseline representation. Early viral response (4 dpi) triggered a visible remodelling of the UMAP landscape. There was an emergence and expansion of specialized activation states, notably the interferon-responsive (G5b, ISG^+^) and Immunosuppressive (G5a) populations. The progression from day 4 to day 8 reveals distinct shifts in both cell population dynamics and functional capacity. By the eighth day, the body mobilized a large, dense cluster of immature neutrophils (G3), alongside the continued growth of the inflammatory G5c group, which indicates the immune system was actively recruiting and maturing new cells to sustain its fight against the ongoing infection. The clusters corresponding to activated/inflammatory (G5c) and Immature neutrophil subsets (G3) neutrophils exhibit prominent expansion, reflecting a sustained systemic and localized immune response to Dengue virus progression (Fig. 3C).

Consistent with our previous report (Duggal et al., 2023), flow cytometric analysis of bone marrow cells isolated from the femurs and tibias of DV-2-infected mice at 4 dpi showed a significant expansion of the CD11b^+^Ly6C^int^Ly6G^low^ (Ly6G^low^) cell population (Fig. 3D). In contrast, the CD11b^+^Ly6C^int^Ly6G^hi^ mature cells (Ly6G^hi^) remained stable (Fig. 3D, right panel). Together, these data identify the emergence of Ly6G^low^ neutrophil population in the BM following DV-2 infection, further validating our scRNA-seq data.

We next checked the presence of this population in the blood and spleen of the infected mice. Interestingly, observed that the proportion of these Ly6G^low^ cells was significantly elevated in the spleen and peripheral blood of DV-2-infected mice (Fig. 3E, F). This builds on the fact that Ly6G^low^ is not only produced in response to infection but also systemically mobilized during infection, suggesting its potential involvement in immunomodulation and disease pathogenesis. Integrating these *in vivo* findings with the scRNA-seq analysis, G5c neutrophil cluster may represent the transcriptional counterpart of the Ly6G^low^ population.

### The Ly6G^low^ neutrophil population significantly differ from the Ly6G^hi^ population in the dengue-infected bone marrow of mice

To gain more insights into Ly6G^low^ neutrophils, we checked the morphology of sorted Ly6G^low^ neutrophils. Giemsa staining and confocal microscopy revealed distinct nuclear morphologies, with Ly6G^low^ neutrophils exhibiting a ring-shaped nucleus, in contrast to the segmented nuclei of their Ly6G^high^ counterparts (Fig. 4A, B). To further confirm that Ly6G^low^ cells are metabolically distinct, the glucose uptake assay showed that they had the highest glucose uptake (Fig. 4C, top, left panel), suggesting increased metabolic demand during infection. Infection significantly increases lipid accumulation in both Ly6G^hi^ and Ly6G^low^ (Fig. 4C, top, right panel), likely indicating the formation of lipid droplets, a common feature of inflammatory responses.

**Fig. 4.**
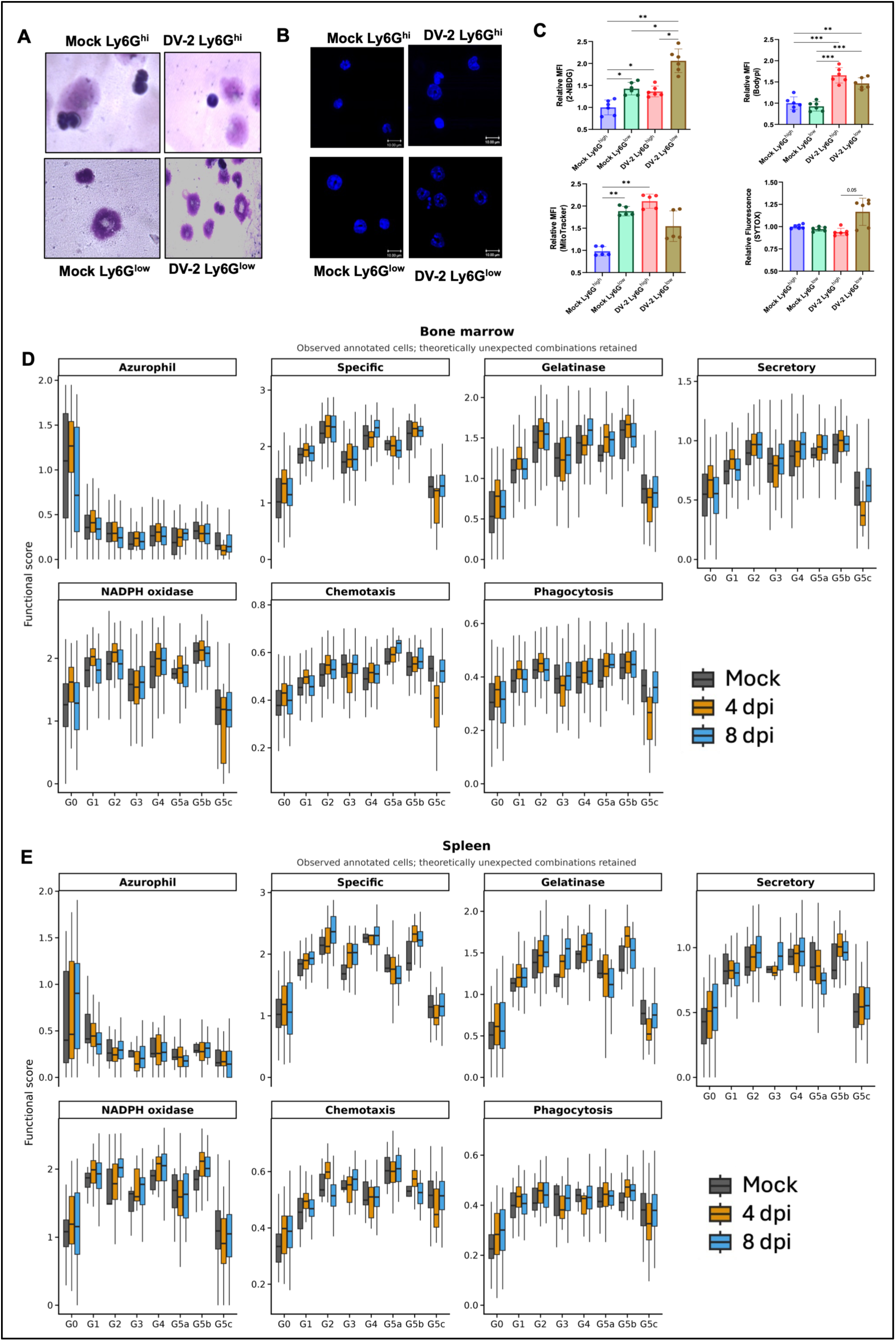
Phenotypic, metabolic, and functional dynamics of neutrophil subsets during DV-2 infection. **(A)** Representative light microscopy images showing the nuclear morphology of Giemsa- or Wright-stained sorted Mock and Dengue virus 2 (DV-2) infected Ly6G^hi^ and Ly6G^low^ neutrophil populations. **(B)** Representative fluorescence microscopy images showing nuclear nuclear lobulation/staining (DAPI/Hoechst) in Mock and DV-2 treated Ly6G^hi^ and Ly6G^low^ neutrophil subsets. Scale bars = 10 *μ*M. **(C)** Flow cytometric quantification of metabolic and cellular parameters in sorted neutrophil subsets across treatment conditions: relative mean fluorescence intensity (MFI) of glucose uptake (2-NBDG, top-left), lipid content (Bodipy, top-right), mitochondrial mass/potential (MitoTracker, bottom-left), and cell death/membrane integrity (SYTOX, bottom-right). Data are presented as mean ± s.d. Statistical significance was determined using one-way ANOVA with Tukey’s post-hoc test; *P < 0.05, P < 0.01, ***P < 0.001. **(D, E)** Module score dynamics reflecting functional pathways (Azurophil, Specific, Gelatinase, Secretory granule proteins, NADPH oxidase activity, Chemotaxis, and Phagocytosis) across defined neutrophil developmental/functional clusters (G0, G1, G2, G3, G4, G5a, G5b, G5c) in the bone marrow (**D**) and spleen (**E**) during Mock, 4 days post-infection (4 dpi), and 8 dpi with DV-2. Box plots indicate the median (centre line), interquartile range (bounds of box), and minimum/maximum values (whiskers).

In the mock mice, Ly6G^low^ cells naturally maintain higher mitochondrial levels than Ly6G^hi^ cells. DV-2 infection significantly boosts mitochondrial activity/content specifically in the Ly6G^hi^ population, reaching levels similar to or exceeding those of Ly6G^low^ cells, suggesting under infection conditions, both Ly6G^low^ and Ly6G^hi^ neutrophil subsets maintained significantly increased mitochondrial activity (Fig. 4C, bottom, left panel). The Sytox assay confirmed that upon exposure to Dengue virus type 2 (DV-2), Ly6G^low^ neutrophils show significantly higher membrane permeability or extracellular DNA release compared to Ly6G^hi^ neutrophils. This suggests that the Ly6G^low^ subset is preferentially vulnerable to DV-2-induced cell death or is more prone to undergoing NETosis upon experimental PMA stimulation (Fig. 4C, Bottom, right panel). Overall, our data indicate that Ly6G^low^ neutrophils are more metabolically active and responsive during a DV-2 infection than their Ly6G^hi^ counterpart.

Functional scoring profiles showed distinct expression patterns for granule subsets (azurophil, specific, gelatinase, and secretory) and functional pathways (NADPH oxidase, chemotaxis, and phagocytosis) across developmental stages G0 to G5c. Functional scoring within the bone marrow and spleen compartments confirmed that these defined developmental states correspond to specialised effector profiles. While early progenitor states (G0/G1) exhibited high azurophilic granule scores, the functional scores for both NADPH oxidase activity and phagocytosis, along with Gelatinase and Secretory granules, were notably high in the G5a and G5b sub-populations in viral infection, indicating a primed, highly enhanced effector functional state (Fig. 4D, E). However, the scores for G5c during active infection (4 dpi and 8 dpi) remain substantially lower than those of the G4 or G5a/G5b resting pool.

The sharp drop in granule content and metabolic capacity (Specific, Gelatinase, Secretory, NADPH oxidase) in G5c, compared to the precursor and mature stages (G1–G5b), typically reflects terminal maturation and is associated with greater degranulation. While effector capacity (such as ROS generation via NADPH oxidase and enzyme secretion) is markedly down-regulated, basic cellular processes like Chemotaxis and Phagocytosis remain partially preserved, suggesting that G5c cells retain motility and uptake capabilities despite losing active granule production. Overall, G5c represents a functionally depleted or late-stage degranulated cell state that retains basic motility and phagocytosis but has lost active granule production and the oxidative burst. Although G5c lacks effector granular content, transcript analysis confirmed that compared to other mature phenotypes (e.g., G4, G5a, G5b), G5c was characterised by the expression of *Il1b, Ptgs2, Clec7a,* and immediate-early genes such as *Fos* and *Jun*. Lacking cell-cycle (*Spc25*) and developmental (*Olfm4*) markers (Fig. 2E), this cluster appears to be defined by functional activation rather than ontogeny.

### Phenotypic characterisation and functional consequences of Ly6G^low^ neutrophils

Next, we aim to investigate the function of the newly formed Ly6G^low^ population in the BM. Many reports suggest that neutrophils produced during emergency granulopoiesis exhibit a modulated function (33, 34), with the capacity to suppress the immune response. In our scRNA and in vivo data, we also observed a significant decrease in the number of CD4+ T cells in the blood of DV-2-infected mice (Fig. 5A-C). However, an increase in the Treg population was observed over the course of infection (Fig. 5D, E). To determine whether Ly6G^low^ neutrophils modulate T cell responses, Ly6G^low^ and Ly6G^hi^ populations were sorted from bone marrow, and CD4⁺ and CD8⁺ T cells were isolated from spleens. Co-culture assays were performed to assess the impact of these subsets on T cell activation and function. T cells were stimulated with plate-bound anti-CD3, neutrophils were added at a 1:10 ratio, and the mixture was cultured for 48 h. Cells were treated with monensin before intracellular cytokine analysis. Ly6G^low^ neutrophils markedly suppressed T cell effector functions (Fig. 5F).

**Fig. 5.**
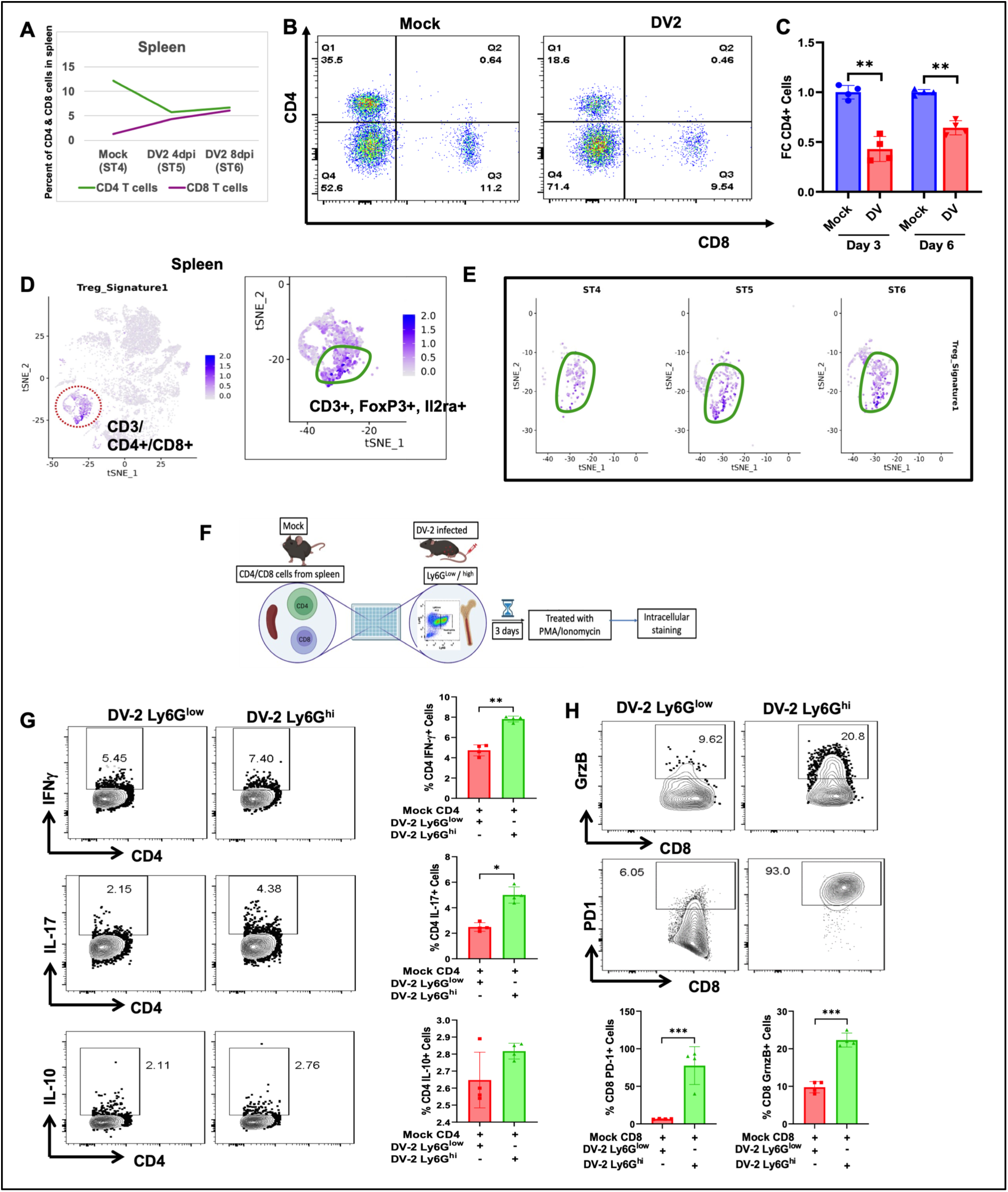
Spleen T cell dynamics and functional modulation during DV-2 infection. (**A)** Quantitative Analysis of Splenic T Cell Percentages. Line graph tracking the percentage of CD4^+^ and CD8^+^ T cells in the spleen from Mock through 4 dpi and 8 dpi. **(B)** Flow Cytometric Validation of CD4/CD8 Frequency. Representative dot plots showing CD4 *vs*. CD8 expression in Mock and DV-2 4 dpi spleen samples. **(C)** The bar graph (right) quantifies the fold change (FC) in CD4^+^ cells, showing a significant decrease at Days 3 and 6 post-infection (indicated by ** p < 0.01). **(D)** Identification of Splenic Treg Populations via scRNA-seq. t-SNE visualisation of splenic cell populations highlighting the Treg_Signature1 cluster. The zoomed-in feature plot confirms the cluster as CD3^+^, FoxP3^+^, and Il2ra^+^ (CD25) regulatory T cells. **(E)** Comparative t-SNE plots showing the density and distribution of the Treg signature in the spleen at different time points: Mock (ST4), 4 days post-infection (ST5), and 8 days post-infection (ST6). **(F)** Functional Impact of Ly6G^hi^ vs. Ly6G^low^ Cells on T Cell Activation. Experimental Schematic: CD4^+^ and CD8^+^ T cells isolated from the spleen were co-cultured with Ly6G^low^ or Ly6G^hi^ cells from DV-2 infected mice for 3 days, followed by PMA/Ionomycin stimulation and intracellular staining. **(G)** CD4^+^ T Cell Cytokine Profile: Intracellular staining for IFN-γ, IL-17, and IL-10 in CD4^+^ T cells. Co-culture with Ly6G^hi^ cells significantly enhances the frequency of IFN-γ+ and IL-17^+^ CD4^+^ T cells (** p < 0.01). **(H)** Representative plots and bar graphs show a significant increase in the percentage of Granzyme B (GrzB)^+^ and PD-1^+^ CD8^+^ T cells when co-cultured with Ly6G^hi^ cells compared to Ly6Glow or Mock (*** p < 0.001).

CD4⁺ T cells co-cultured with Ly6G^low^ neutrophils exhibited significantly decreased frequencies of IFN-γ⁺ (Th1) and IL-17⁺ (Th17) cells (Fig. 5G). Similarly, in CD8⁺ T cells, co-culture with Ly6G^low^ cells resulted in reduced granzyme B compared to Ly6G^hi^ controls (Fig. 5H). Collectively, these findings demonstrate that Ly6G^low^ neutrophils exert potent immunosuppressive effects on both CD4⁺ and CD8⁺ T cell responses.

### Dengue infection augmented FGL2 release from the Ly6G^low^ neutrophil population and suppressed T cell function

Given the distinct appearance of Ly6G^low^ neutrophils upon infection and the suppression of T cell function, we examined the secretome of Ly6G^low^ cells. We observed from our scRNA data that only G5c neutrophils showed a significant increase at the transcript level with Fibrinogen-like protein-2 (FGL2) expression (Log2Fold change = 3.34) (Fig. 6A) and correlated with Ly6G expression. FGL2 encompasses mFGL2 (membrane-bound) and sFGL2 (soluble), which differ in tertiary structure; these differences underlie their functional differences. sFGL2 only contains the C-terminal FRED domain, which underlies its immunomodulatory effects. To confirm that FGL2 is secreted, and increased FGL2 levels are more specific to dengue infection and not an outcome of any systemic infection, we infected AG129 mice with Japanese encephalitis virus (JEV) and dengue virus and measured FGL2 levels in serum during severe symptomatic conditions. Our data confirmed that infection with flaviviruses (JEV and DV-2) upregulates serum FGL2 expression in AG129 mice. But DV-2 infection induces a significantly stronger FGL2 response compared to JEV infection. we measured FGL2 levels in the serum of infected mice (Fig. 7C). Further study confirmed that FGL2 serum levels are significantly upregulated during Dengue virus (DV-2) infection in AG129 murine models, both at 4 and 8 dpi (Fig. 6D) and human patients, with human Dengue-positive individuals showing the highest overall elevations compared to healthy controls and other febrile illness (OFI) patients (Fig. 6E). Compared with healthy donor serum, FGL2 levels increased significantly in patients with other febrile illnesses (OFI). However, sera from Dengue patients exhibited significantly higher levels (at least 2-fold) than those from OFI patients (Fig. 6E). We also investigated whether the dengue virus can induce FGL2 secretion from neutrophils. We purified neutrophils from the blood of healthy donors and cultured them with the dengue virus. FGL2 was measured in supernatants collected at 12 and 24 hpi. As shown in Fig. 6F, incubation with dengue virus significantly increased FGL2 levels in the supernatant, confirming that neutrophils secrete sFGL2 in response to DV-2 infection.

**Fig. 6.**
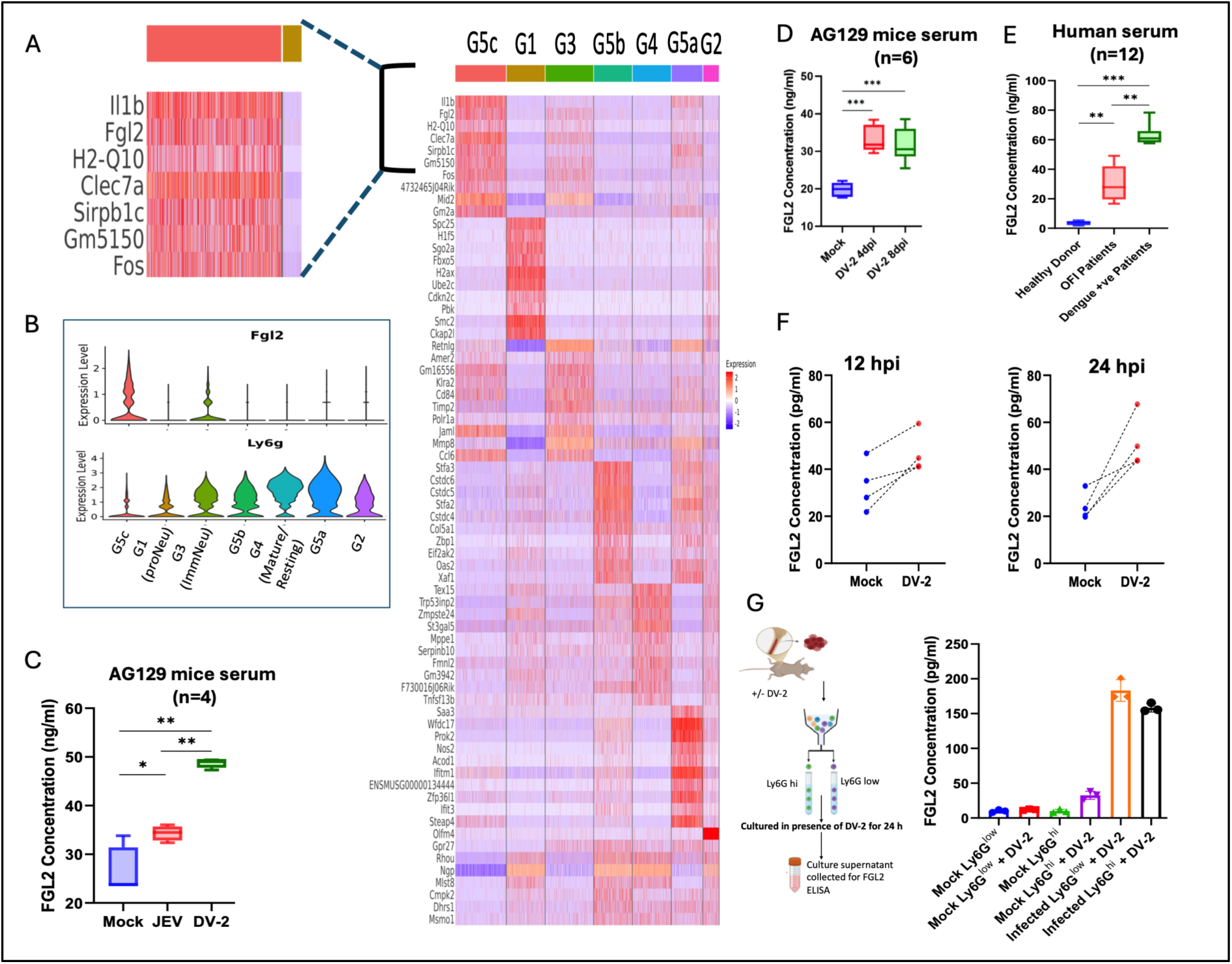
FGL2 expression is upregulated in specific neutrophil subsets and elevated during flavivirus infection. **(A)** Heatmap showing marker gene expression across identified neutrophil clusters (G5c, G1, G3, G5b, G4, G5a, G2). The zoomed-in section highlights high expression of *Fgl2*, *Il1b*, *H2-Q10*, *Clec7a*, *Sirpb1c*, *Gm5150*, and *Fos* within specific subsets. Color bar indicates relative expression levels (z-score log-transformed values). **(B)** Violin plots illustrating single-cell gene expression levels of *Fgl2* and *Ly6g* across neutrophil developmental and functional subsets (G5c, G1/proNeu, G3/ImmNeu, G5b, G4/Mature/Resting, G5a, and G2). **(C)** Quantification of serum FGL2 concentrations (ng/ml) measured by ELISA in AG129 mice following Mock, Japanese Encephalitis Virus (JEV), or Dengue Virus 2 (DV-2) infection (n = 4 mice per group). **(D)** Serum FGL2 concentrations (ng/ml) in AG129 mice at 4 and 8 days post-infection (dpi) with DV-2 compared to Mock-infected controls (n = 6 mice per group). **(E)** Serum FGL2 concentrations (ng/ml) in healthy donors, patients with Other Febrile Illnesses (OFI), and Dengue-positive (+ve) patients (n = 12 individuals per group). **(F)** Pairwise comparisons of FGL2 concentration (pg/ml) from Mock and DV-2 infected samples at 12 hours post-infection (hpi, left) and 24 hpi (right). **(G)** Schematic representation of experimental workflow (left) and quantification of FGL2 concentration in culture supernatants (right) from sorted Ly6G^low^ and Ly6G^hi^ neutrophil populations isolated from Mock or DV-2 infected mice after 24 hours in culture. Data in the box plots (**C**–**E**) show the median, interquartile range (box bounds), and minimum/maximum values (whiskers). Data in bar graphs (**G**) represent mean ± s.d. Statistical significance was determined using two-sided Student’s t-test or one-way ANOVA with Tukey’s post-hoc test; *P < 0.05, P < 0.01, ***P < 0.001.

**Fig. 7.**
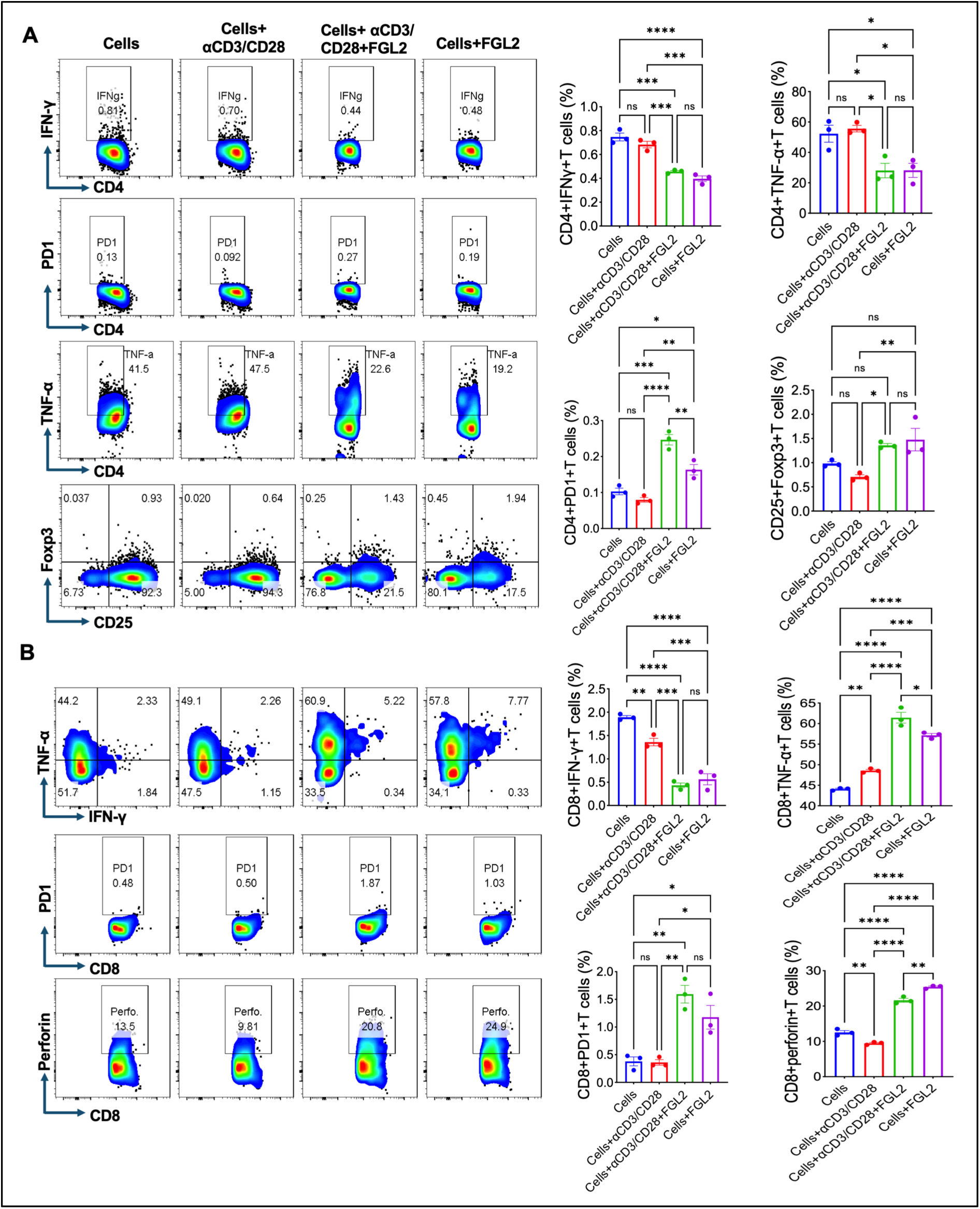
FGL2 suppresses T cell functional activity and promotes an immunosuppressive phenotype. **(A)** Splenocytes were stimulated with anti-CD3 ± anti-CD28 in the presence or absence of recombinant FGL2 or FGL2 alone. Flow cytometric analysis revealed reduced frequencies of CD4⁺IFN-γ⁺ and CD4⁺TNF-α⁺ effector T cells, alongside increased CD4⁺PD-1⁺ and CD25⁺Foxp3⁺ regulatory T cells following FGL2 treatment, indicating suppressed activation and enhanced inhibitory signalling (n=3). **(B)** Intracellular analysis of CD8⁺ T cells demonstrated reduced IFN-γ, TNF-α, and perforin production, with increased PD-1 expression in the presence of FGL2, consistent with an immunosuppressive/exhausted phenotype. Data are representative of independent experiments(n=3). *p < 0.05, ** p < 0.01, *** p < 0.001, **** p < 0.0001.

To further confirm that Ly6G^low^ neutrophils are secreting a high amount of FGL2, we purified Ly6G^low^ neutrophils from the bone marrow of mock and infected mice. We stimulated them in the presence and absence of the dengue virus. Since we observed Ly6G^low^ neutrophils released into the blood and spleen of infected mice, we hypothesized that these primed neutrophils, upon interaction with DV-2 in circulation, could release higher levels of sFGL2. To mimic this situation, we incubated Ly6G^low^ and Ly6G^hi^ cells purified from mock and infected mice’s bone marrow (Fig. 6G, left panel). As shown in Fig. 6G (right panel), mock Ly6G^low^ did not release FGL2 into the supernatant, even in the presence of dengue virus; however, Ly6G^hi^ from mock mice in the presence of dengue virus released higher levels of FGL2 into the supernatant. This observation is similar to our previous results (Fig. 6E), in which Ly6G^hi^ neutrophils in the presence of virus augmented sFGL2 release.

Interestingly, we observed that Ly6G^low^ or Ly6G^hi^ purified from the bone marrow of infected mice showed 3-4-fold higher FGL2 expression in the supernatant compared to mock Ly6G^hi^ neutrophils (Fig. 6G). All these data confirmed that dengue virus infection augmented FGL2 release from activated neutrophils, and Ly6G^low^ neutrophils were the major source of FGL2.

FGL2 plays a multifaceted role in the immune system and acts as an important immunosuppressive effector cytokine of Tregs, suppressing Th1 cytokine production while enhancing Th2 cytokine production (35, 36). Given the immunosuppressive activity of Ly6G^low^ neutrophils, we next examined whether soluble FGL2 (sFGL2) directly modulates T cell responses. Splenocytes were stimulated with anti-CD3/CD28 in the presence or absence of recombinant FGL2 (1 µg ml⁻¹), followed by flow cytometric analysis of T cell activation and functional markers (Fig. 7A, B). FGL2 treatment significantly reduced the frequencies of CD4⁺IFN-γ⁺ and CD4⁺TNF-α⁺ T cells compared to activated controls, indicating suppression of effector responses. This was accompanied by increased PD-1 expression and an expansion of CD25⁺Foxp3⁺ regulatory T cells, consistent with enhanced inhibitory signalling and regulatory skewing. Further, we investigated how the protein FGL2 (Fibrinogen-like protein 2) affects the function and exhaustion of CD8+ T cells. In CD8⁺ T cells, FGL2 reduced IFN-γ production while increasing PD-1 expression, indicative of an exhaustion-like phenotype. Notably, TNF-α and perforin expression remained increased in CD8+ T cells, suggesting partial retention of effector function despite functional impairment. Together, these findings indicate that FGL2 does not induce complete T cell suppression but rather drives a state of dysfunctional activation characterized by impaired cytokine production and heightened inhibitory signaling. Given that DV-2–exposed neutrophils secrete soluble FGL2 (sFGL2) (Fig. 6G), we next assessed the functional impact of neutrophil-derived secretome on T cell responses. Splenocytes were stimulated with conditioned supernatants derived from Ly6G^hi^ and Ly6G^low^ neutrophils isolated from mock- and DV-2–infected mice (Fig. 8A– D).

**Fig. 8.**
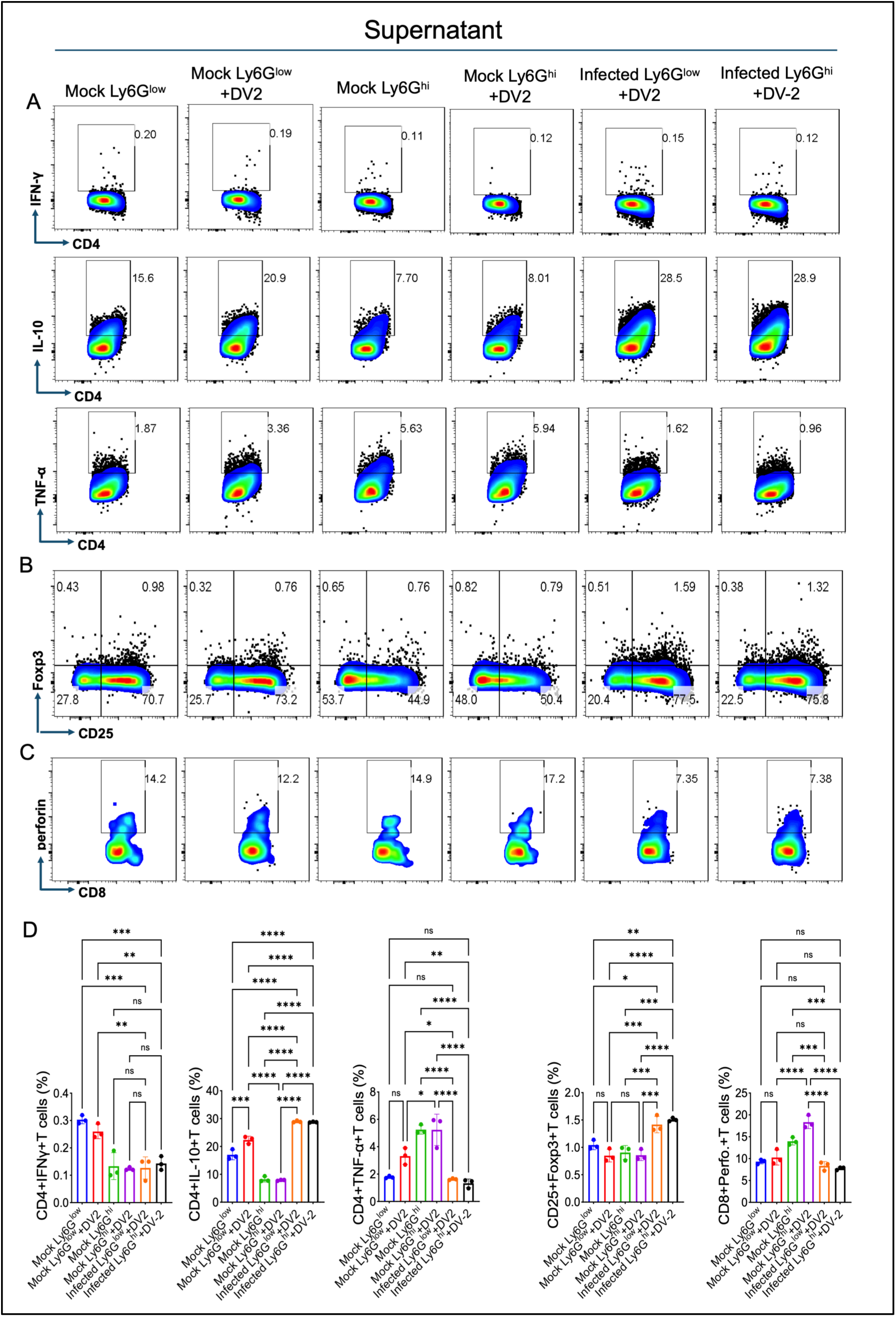
Neutrophil-derived FGL2-containing supernatant suppresses T cell responses during dengue infection. Conditioned media were generated from Ly6G^low^ and Ly6G^hi^ neutrophils isolated from mock- and DV-2–infected mice and used to stimulate draining lymph node cells. (**A)** Intracellular cytokine analysis revealed reduced IFN-γ and TNF-α production, accompanied by increased PD-1 expression and IL-10 levels in T cells stimulated with DV-2-infected Ly6G^low^ and high neutrophil-conditioned media, indicating suppressed effector function and enhanced inhibitory signalling. **(B-D**) Flow cytometric analysis showed increased frequencies of CD25⁺Foxp3⁺ regulatory T cells and reduced CD8⁺perforin⁺ effector T cells following stimulation with DV-2-infected Ly6G^low^ and high neutrophil-conditioned media. Data are representative of independent experiments. *p < 0.05, ** p < 0.01, *** p < 0.001, **** p < 0.001.

Supernatants from DV-2–primed neutrophils, which exhibited elevated sFGL2 levels, markedly suppressed CD4⁺ T cell effector responses, as evidenced by reduced frequencies of IFN-γ⁺ and TNF-α⁺ cells, alongside increased IL-10⁺ CD4⁺ T cells and expansion of CD25⁺Foxp3⁺ regulatory T cells. These findings are consistent with an immunosuppressive and regulatory skewing of CD4⁺ T cell responses. In contrast, conditioned media from DV-2–primed neutrophils did not enhance perforin expression in CD8⁺ T cells compared to mock controls, indicating impaired cytotoxic function. This hints at the possible contribution of other neutrophil-derived factors to T cell responses; however, the role of FGL2 cannot be overlooked.

Overall, FGL2 acts as a key immunosuppressive mediator, blunting CD4⁺ and CD8⁺ T cell effector cytokine production IFN-γ⁺, driving T cell exhaustion (PD-1 upregulation), and promoting regulatory T cell (CD25⁺Foxp3⁺) expansion. DV-2-infected neutrophil secretome further drives systemic immune dysfunction by promoting IL-10 release and Treg expansion while impairing pro-inflammatory effector cytokine responses and cytotoxic capabilities.

## Discussion

This study outlines a dynamic shift in the neutrophil landscape in which the DV-2 hijacks the body’s natural defence mechanisms, turning an emergency immune response into immunosuppression. To elucidate neutrophil dynamics in dengue and translate single-cell insights into clinical applications, this study included three crucial components: (i) a representative disease model reflecting DV-induced neutrophil heterogeneity, (ii) optimised sampling kinetics that reflect the dynamics of neutrophil heterogeneity, mild and severe conditions, and (iii) robust *ex vivo and in vitro* models to characterise the functional profiles of the identified neutrophil subsets.

At the hematopoietic level, the initial response to DV-2 in AG129 mice is a rapid, systemic increase in the production of interferon-responsive granulocytes (G5b) and immunosuppressive neutrophil (G5a) in the bone marrow. AG129 mice are Type-I and II interferon receptor-deficient; however, interferon lambda receptors are functional. The G5b neutrophils carry a high expression of interferon lambda. The early response of G5b is likely mediated by IFN-lambda. The appearance of both G5a and G5b indicates a balanced environment in which host immune systems sought to maintain an antiviral state while suppressing excessive pro-inflammatory responses.

While neutrophils usually play a protective role in response to infection, our data suggest that in Dengue, this process is deviated in severe conditions. Instead of producing mature neutrophils, the bone marrow undergoes a profound reorganisation, shifting its output toward specialised, pro-inflammatory, and immunosuppressive Ly6G^low^ subsets as well as immature subsets, associated with severe dengue.

Based on the shared transcriptome signature and appearance in UMAP, Ly6G^low^ neutrophils appear to be closer to G3 immature neutrophils. Under severe conditions, G3 immature neutrophils probably converted to the G5c Ly6G^low^ population, which expanded in the bone marrow. The Ly6G^low^ population produced in the BM is detectable in the blood and spleen, thereby highlighting its possible role in immunopathogenesis. Another critical finding is the augmentation of Fibrinogen-like protein 2 (FGL2), also known as fibroleukin. FGL2 is elevated both within the Ly6G^low^ neutrophils and as a soluble protein in the serum of mice and humans during viral infections. FGL2 promotes pathogenesis through two structural forms: membrane-bound FGL2 (mFGL2), which acts as a prothrombinase, inducing microvascular thrombosis and tissue necrosis in HBV- and HCV-infected patients, and soluble FGL2 (sFGL2), which functions as an immunosuppressor. These mechanisms contribute to the chronic immune dysfunction observed in HIV and other viral pathologies (37–41).

Soluble FGL2 (sFGL2) binds to the inhibitory receptor FcgammaRIIB on various immune cells (42). In many infections, elevated FGL2 levels are a strong biomarker for disease severity and poor prognosis. High sFGL2 levels correlate with high viral loads and liver failure. In mouse models (MHV-3), blocking FGL2 or using *fgl2*-deficient mice prevents lethal hepatic necrosis. FGL2 is upregulated in patients with severe respiratory distress. Specifically in COVID-19, it has been linked to the depletion of CD8+ T cells via the FcgammaRIIB-mediated apoptotic pathway (42, 43).

In our study, G5c (Ly6G^low^) was the only cell type to express high levels of Fgl2. Upon interacting with DV-2, these neutrophils significantly increased their release of soluble FGL2 (sFGL2). This sFGL2-rich supernatant directly impairs T cell function by: (1) reducing effector cytokine production (e.g., IFN-gamma); increasing exhaustion markers, and inducing a suppressive, regulatory phenotype. While a lack of knockout models and neutralizing antibodies prevented us from confirming its role in inflammation enhancement, other research links FGL2 to procoagulant and immune-regulatory responses. It was reported earlier that mFGL2 activates thrombin, establishes a feedback loop with C5a and TNF-α, and simultaneously enhances NET formation through the FGL2-MCOLN3-autophagy axis (41).

In dengue infection, NETosis plays a vital role in disease severity (16, 44, 45). Future studies in FGL2 knockout mice may be needed to confirm its role in immune inflammation. Importantly, FGL2 levels were significantly upregulated in the plasma of dengue-infected patients, suggesting the need for larger, more diverse cohorts of dengue patients to evaluate it as a biomarker of dengue severity.

Taken together, our study concludes that (1) DV-2 infection triggers expansion of hematopoietic progenitor cells in the bone marrow. (2) Infection reorganises the myeloid compartment, shifting bone marrow production toward a specialized, immunosuppressive, and proinflammatory Ly6G^low^ neutrophil subset. (3) Ly6G^low^ neutrophil secretes Fibrinogen-like protein 2 (FGL2) and drives T cell exhaustion (Fig. 9). Thus, our study confirms that DV-2 is associated with specialised neutrophil subsets to drive FGL2-mediated immunosuppression and is associated with dengue disease progression in the mouse model.

**Fig. 9.**
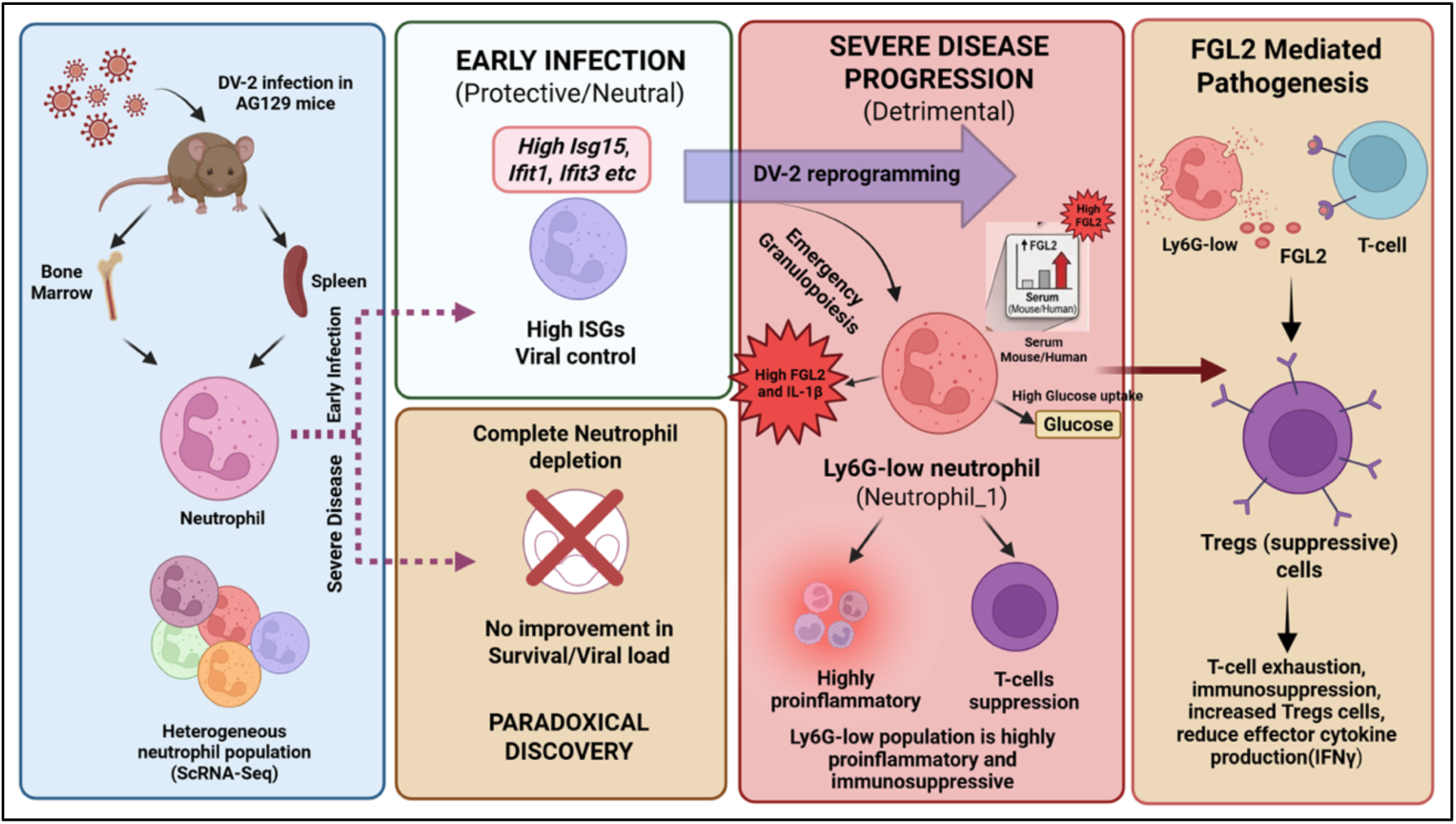
Schematic overview showcasing the progression of Dengue Virus 2 (DV-2) infection in an AG129 mouse model, transitioning from an early protective phase to a detrimental, immunosuppressive severe disease state.

## Contributors

**SR:** Conceptualisation, ScRNA Library preparation, Formal analysis, Data curation, Investigation, Animal experiments, Methodology, Writing and Editing – original draft. **AC, SN:** In vitro studies, Investigation, Animal experiments, Functional analysis, Assisting SCRNA-Sequencing and Data analysis. **ADP:** ScRNA-Seq quality check, ScRNA-Seq analysis pipeline development, data curation and analysis, data submission**. SS, AA:** Resources, T cells functional study, data acquisition, data analysis, experimental set-up, Methodology, Writing and Editing – original draft. **GS**, **SwS:** animal experiments, virus resources, Editing – original draft. **AB:** Conceptualisation, Resources, Funding, Supervision, Writing and editing – original draft.

## Acknowledgements

We sincerely acknowledge Dr Romsha Kumar, Sr. Application Scientist, Single Cell MultiOmics, BD Life Sciences-BioSciences, India/South Asia, for her technical support in designing the experimental setup and support for the smooth execution of ScRNA-sequencing. We thank the Experimental Animal Facility (EAF) at the NCR-Biotech Science Cluster for providing infrastructure for small-animal experiments. We sincerely thank the EAF staff for their assistance in ensuring the study’s smooth conduct. The graphical abstract was *created in BioRende*r.

